# The fitness cost and benefit of phase separated protein deposits

**DOI:** 10.1101/574657

**Authors:** Natalia Sanchez de Groot, Marc Torrent Burgas, Charles N. J. Ravarani, Ala Trusina, Salvador Ventura, M. Madan Babu

## Abstract

Phase separation of soluble proteins into insoluble deposits is associated with numerous diseases. However, protein deposits can also function as membrane-less compartments for many cellular processes. What are the fitness costs and benefits of forming such deposits in different conditions? Using a model protein that phase separates into deposits, we distinguish and quantify the fitness contribution due to the loss or gain of protein function and deposit formation in yeast. The environmental condition and the cellular demand for the protein function emerge as key determinants of fitness. Protein deposit formation can lead to cell-to-cell differences in free protein abundance between individuals. This results in variable manifestation of protein function and a continuous range of phenotypes in a cell population, favoring survival of some individuals in certain environments. Thus, protein deposit formation by phase separation might be a mechanism to sense protein concentration in cells and to generate phenotypic variability. The selectable phenotypic variability, previously described for prions, could be a general property of proteins that can form phase separated assemblies and may influence cell fitness.

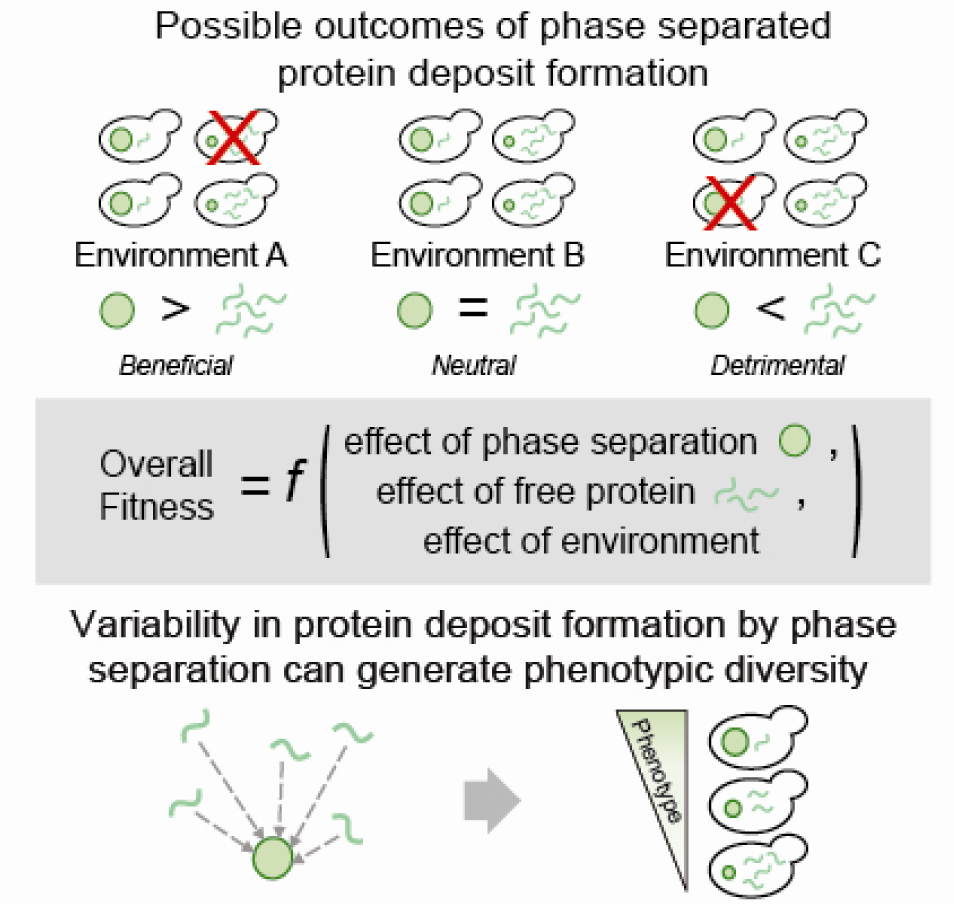

**Stand-first text:** Using a model protein that phase separates into deposits, we distinguish and quantify the fitness contribution due to the loss or gain of protein function and deposit formation in yeast.

**Bullet points:**

- The presented approach identifies and quantifies different fitness effects associated with protein deposit formation due to phase separation
- The environmental condition and the cellular demand for the protein function emerge as key determinants of fitness upon protein deposit formation
- Variability in protein deposit formation can lead to cell-to-cell differences in free protein abundance between individuals
- Protein phase separation can generate a continuous range of phenotypes in a cell population

## INTRODUCTION

The exposure of certain polypeptide segments in a protein to the solvent can trigger a process in which proteins phase separate into macromolecular assemblies (Veis, 2011). The formation of protein deposits can affect biological processes as a result of a loss and/or gain of function, and have been implicated in disorders such as Alzheimer’s or Parkinson’s disease (Babu et al, 2011; Chiti & Dobson, 2006; Ciryam et al, 2013; Gsponer & Babu, 2012; Jahn & Radford, 2008; Lin et al, 2015; Sanchez de Groot et al, 2012; Woerner et al, 2016). However, protein sequences predisposed to form deposits are found in all kingdoms of life suggesting a neutral or advantageous effect on cell fitness (Berchowitz et al, 2015; Chavali et al, 2017a; Khan et al, 2015; Li et al, 2012; Newby & Lindquist, 2013). In fact, it has been shown that phase separation promoting sequences are essential to build membrane-less structures and higher-order assemblies with several biological functions (Bershtein et al, 2012; Chavali et al, 2017a; Ciryam et al, 2013; Gsponer & Babu, 2012; Holehouse & Pappu, 2018; Lin et al, 2015; Maji et al, 2009; Miller et al, 2015; Nott et al, 2015; Sanchez de Groot et al, 2012; Suresh et al, 2015; Toretsky & Wright, 2014; Wallace et al, 2015; Xiang et al, 2015; Zhu & Brangwynne, 2015).

Protein deposit formation due to phase separation inside the cell is a complex process that depends on a number of features such as the physicochemical properties of the polypeptide sequence (e.g. hydrophobicity, net charge, etc.) (de Groot et al, 2006), local protein concentration (e.g. high vs. low concentration) (Ciryam et al, 2013; Levy et al, 2014; Stepanenko et al, 2016) and its interaction with cellular components (e.g. chaperones, RNA, etc.) (De Baets et al, 2011; Gsponer & Babu, 2012; Jain et al, 2016; Maharana et al, 2018; Miller et al, 2015; Pak et al, 2016; Sanchez de Groot et al, 2012; Zhang et al, 2015). The different combinations of these features result in the formation of deposits with different physicochemical and dynamic properties (**Figure 1A**). According to their viscoelastic characteristics and their ability to exchange components with the cytoplasm, protein assemblies can adopt a wide range of states that go from highly dynamic states with liquid droplet properties to almost static states with solid-like properties (**Figure 1A**) (Franzmann et al, 2018). In this manner, phase-separated assemblies can be classified as liquid-liquid (e.g. nucleolus) or liquid-solid (e.g. amyloid aggregate) according to their viscoelastic and dynamic properties (e.g. deformation and coalescence) (Alberti et al, 2019; Boeynaems et al, 2018; Escusa-Toret et al, 2013; Franzmann et al, 2018; Jahn & Radford, 2008; Jain et al, 2016; Kaganovich et al, 2008; Lin et al, 2015; Miller et al, 2015; Morley et al, 2002; Toretsky & Wright, 2014; Zhang et al, 2015; Zhu & Brangwynne, 2015).

**Figure 1.**
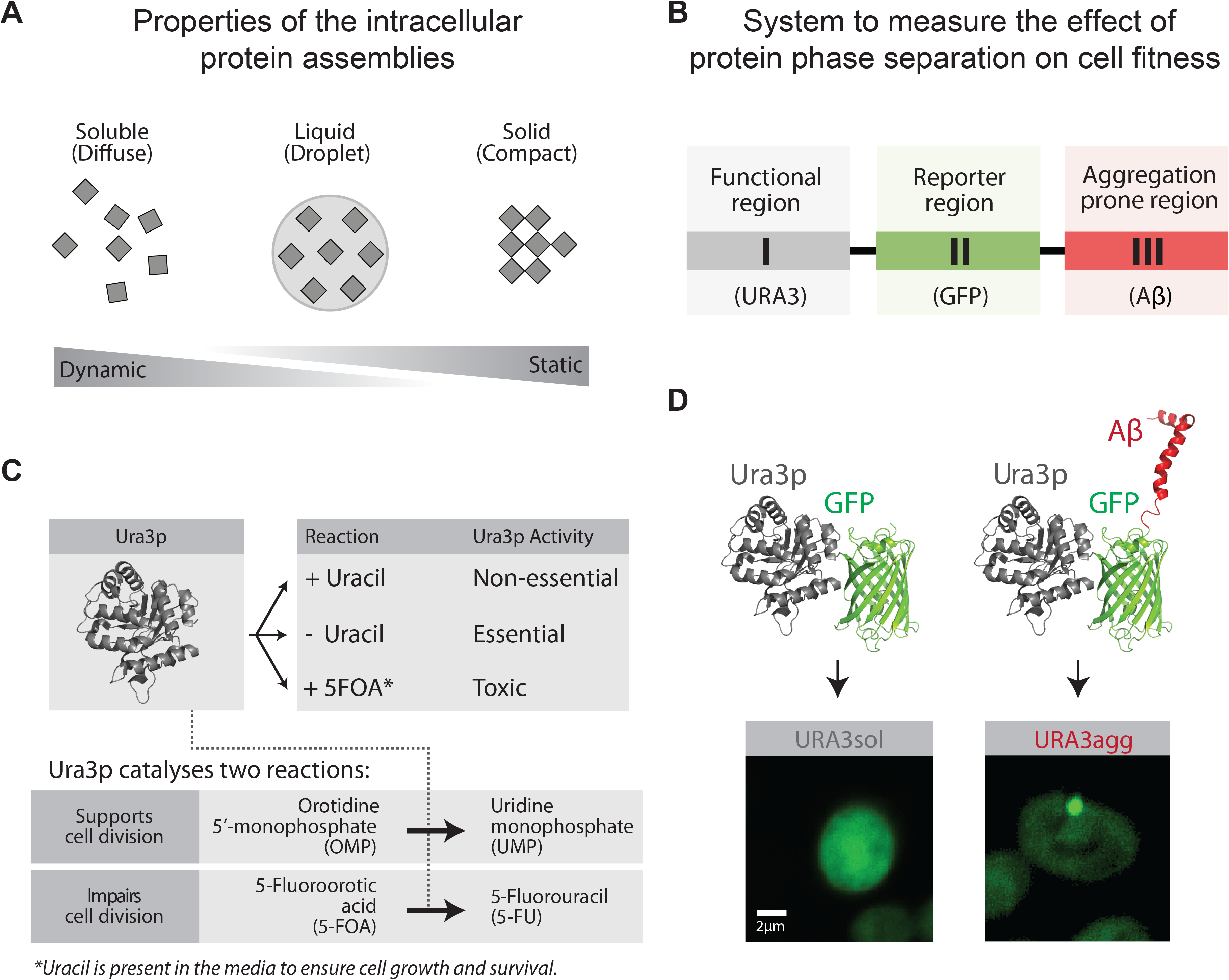
Model proteins and design of experimental system. **A.** Protein phase separation can lead to the formation of protein assemblies with varying dynamics. Soluble proteins may phase separate into liquid-like droplets or insoluble deposits that according to their viscoelastic properties and their ability to exchange components with the cytoplasm vary from being dynamic to static. **B.** Design of the modular system to measure the effect of protein phase separation on cell fitness. **C.** Growth media composition and the essential/non-essential/toxic roles of Ura3p. **D.** The model proteins consist of a fusion between Ura3p and GFP (URA3_sol_, left) and the amyloid-β-peptide of 42 residues (Aβ) (URA3_agg_, right). After 18 hours of expression Ura3p_sol_ remains homogeneously distributed whereas Ura3p_agg_ is accumulated into intracellular foci.

This variety of assemblies with distinct dynamic behavior (**Figure 1A**) is employed by cells to regulate key biological processes such as gametogenesis (Berchowitz et al, 2015), programmed necrosis (Li et al, 2012), hormone storage (Maji et al, 2009), or memory maintenance (Khan et al, 2015), among others functions (Chakrabortee et al, 2016; Miller et al, 2015; Mitrea & Kriwacki, 2016; Toretsky & Wright, 2014). Proteins that are able to phase separate can perform their biological functions in the soluble state such as the pituitary hormones (Maji et al, 2009), the phase-separated state, such as the amyloid-like form of Rim4 (Berchowitz et al, 2015) or in both states (e.g. monomeric Orb2 represses, whereas oligomeric Orb2 activates translation (Khan et al, 2015). Therefore, the formation of phase-separated structures is not only associated with diseases but can positively influence cell fitness under different conditions. For instance, the recruitment of certain metabolic enzymes into cytoplasmic reservoirs/deposits enhances yeast survival during periods of starvation and stress (Franzmann et al, 2018; O’Connell et al, 2014; Petrovska et al, 2014; Riback et al, 2017; Suresh et al, 2015; Wallace et al, 2015). Thus, beyond prion domain containing proteins, the extensive number of reported self-assembly events suggest that this is likely to be an intrinsic property of the polypeptide chain (Dobson, 1999; Monsellier et al, 2008). The ability for proteins to phase separate tend to be encoded in specific regions or domains (**Appendix Figure S1**) (Berchowitz et al, 2015; Hervas et al, 2016; Khan et al, 2015; Li et al, 2012; Marsh & Teichmann, 2010; Moore et al, 2008). Such a modular organization permits independent evolution of protein regions and facilitates the emergence of proteins with new properties through recombination (Marsh & Teichmann, 2010; Moore et al, 2008).

The ultimate fitness consequences of phase separated structures will depend on the function of the protein, the components sequestered in the assembly, dynamic nature and the molecular structure of the phase separated assembly among other factors. Due to the complexity and the diverse effects associated with phase separation, there is a need to develop rational approaches that permit the quantification of the effects of forming such deposits on cell fitness in multiple environments and determine how proteins with phase separation prone segments are selected for, or against, in a cell population. To begin unravelling the effect of these factors, we have designed a model protein that phase separates to primarily form insoluble deposits and describe a population genetics approach. This allows us to disentangle and quantify the effect of the following factors associated with phase separation in different environments: the fitness change associated with (i) deposit formation, (ii) the loss of an essential biochemical activity of the protein that forms the deposit, and (iii) the gain of a beneficial effect due to deposit formation.

## RESULTS

### Model for protein phase separation: One protein three roles

Various studies have reported diverse, and often conflicting effects in terms of the beneficial and detrimental effects of protein deposit formation on cell fitness (Escusa-Toret et al, 2013; Geiler-Samerotte et al, 2011; Maji et al, 2009; Sanchez de Groot et al, 2012; Tomala et al, 2014). These differences are understandable if one considers the complexity of deposit formation, the differences in the dynamic nature of the deposit, biochemical function of the protein forming the deposit, and the diversity in the experimental conditions in which the studies have been carried out. Furthermore, prior studies in the literature typically do not explicitly discriminate the different phase separation processes or the different fitness effects associated with proteins (e.g. loss vs. gain of function due to deposit formation). Here, we measure and disentangle the effects of phase separation of a model protein. To trigger the phase separation process, instead of mutating an endogenous protein to destabilize it as has been done before (Geiler-Samerotte et al, 2011; Tomala et al, 2014), we designed a modular system that allows us to disentangle and quantify the cost/benefit of protein phase separation while tuning different roles influencing this process (e.g. the essentiality/non-essentiality/toxicity of a protein). The model protein phase separates from a mainly soluble, functionally active state into a primarily insoluble, functionally less active state (**Material and Methods** and **Figure 5**, **Figure EV1**, **Appendix Figure S2**).

Mimicking the domain organization seen in nature (Derkatch et al, 1996; Marsh & Teichmann, 2010; Moore et al, 2008) (**Appendix Figure S1**), our model protein is made up of three modular components (**Figure 1B**; Material and Methods). The first component is a polypeptide segment whose biochemical activity can be essential, non-essential or toxic for the cell. We chose the endogenous yeast enzyme orotidine-5’-phosphate decarboxylase (Ura3p) involved in the production of pyrimidine nucleotides and widely employed for positive and negative selection (Seiple et al, 2006) (**Figure 1C**). Ura3p activity is essential in yeast cells that are grown in the absence of uracil. However, Ura3p activity is non-essential when grown in the presence of uracil. Furthermore, Ura3p activity is toxic in the presence of an alternative substrate named 5-fluoroorotic acid (5FOA), as it leads to the production of a toxic compound (5-fluorouracil, 5FU) leading to cell division arrest and cell death (Seiple et al, 2006) (**Figure 1C**). The second component is a reporter (Green Fluorescent Protein; GFP) that allows monitoring the integrity and the distribution/location of the protein in the cell (**Figure 1B, 1D**). The third component is a phase separation promoting segment that leads to the formation of intracellular, insoluble protein deposits, which in our system is the 42 amino acid amyloid-β-peptide (Aβ) (de Groot & Ventura, 2006; Sanchez de Groot et al, 2015; Villar-Pique & Ventura, 2013) (**Figure 1D**).

We built two different constructs; one encoding Ura3p fused to GFP (to obtain a protein with no or low phase separation potential, Ura3p_sol_) and another that includes the Aβ peptide to promote the phase separation process (Ura3p_agg_) (**Figure 1D**, **Material and Methods**). Instead of using Ura3p_sol_ as a control, we initially considered fusing Ura3p-GFP to a soluble variant of the Aβ42 (Sanchez de Groot et al, 2015; Villar-Pique & Ventura, 2013) (e.g. a non-foci forming variant). However, the different soluble variants are not always completely soluble in yeast, since after fractionation they are still found in the insoluble part and they form deposits in some of the stress environments investigated in the current work (Villar-Pique & Ventura, 2013). Hence, we decided that the addition of a soluble variant of Aβ42 will not be a suitable control for generating the soluble version of Ura3p. We also decided against fusing another soluble protein with a length similar to Aβ42 but with a different sequence because URA3-GFP fusion is already a large protein and the inclusion of a “random” short soluble peptide sequence might not affect the cost significantly.

We integrated the two chimeric genes (Ura3p_sol_ and Ura3p_agg_) into a stable genomic region (TRP1 locus) to ensure steady expression in multiple generations. We included an inducible promoter (GAL1) to control transcription and guarantee expression under different environments (**Material and Methods**, **Appendix Figure S2**). Their integration in the *S. cerevisiae* genome resulted in two strains (URA3_sol_ and URA3_agg_) with the same genomic background and with similar mRNA expression levels (**Figure EV2**). This ensures that the fitness cost of expressing the constructs as gratuitous proteins, will remain similar in the two strains (Dekel & Alon, 2005; Geiler-Samerotte et al, 2011; Kafri et al, 2016; Pena et al, 2010; Plata et al, 2010) and hence minimally influence the measurements of fitness effects (next section). We monitored competitive growth and formation of deposits in different environments by changing the osmotic pressure (1M Sorbitol, 0.5M NaCl), oxidation level (0.5mM H_2_O_2_, 1mM DTT), temperature (37°, 30° and 25°C) and in the presence of a chemical chaperone (0.5M Proline). After 18 hours of induction at standard growth conditions (30°C) in mid-log phase, Ura3p_sol_ remains distributed through the cytoplasm, whereas Ura3p_agg_ forms stable, non-dynamic protein deposits similar to IPODs (insoluble protein deposits) (Kaganovich et al, 2008), as measured by FRAP experiments (**Figure 1D**, **Figure EV3**). Although some of these assemblies may contain other proteins (Rothe et al, 2018), prior findings suggest that the vast majority of them are likely to be Ura3p_agg_ (Morell et al, 2011; Sanchez de Groot et al, 2015).

### Quantifying selection for/against phase separation in different environments

We experimentally determined how the formation of intracellular protein deposits is selected for, or against, in a population when the two strains (URA3_sol_ and URA3_agg_) are grown in competition (**Figure 2A**). We grew mixed cell cultures (1:1 initial proportion) for three days in exponential phase and measured the selection coefficient (*S*) by PCR (**Figure 2A**, **Appendix Table S1, Material and Methods**) (Chevin, 2011; Geiler-Samerotte et al, 2011; Sanchez de Groot et al, 2015). This growth phase will ensure a constant doubling time, a young population, and a low number of aged protein assemblies (Hill et al, 2016). The selection coefficient (*S*) is related to the difference in growth rate between the two strains (**Figure 2B**) and quantifies how much the cell fitness increases (positive values) or decreases (negative values) due to the formation of deposits in URA3_agg_ in comparison to URA3_sol_.

**Figure 2.**
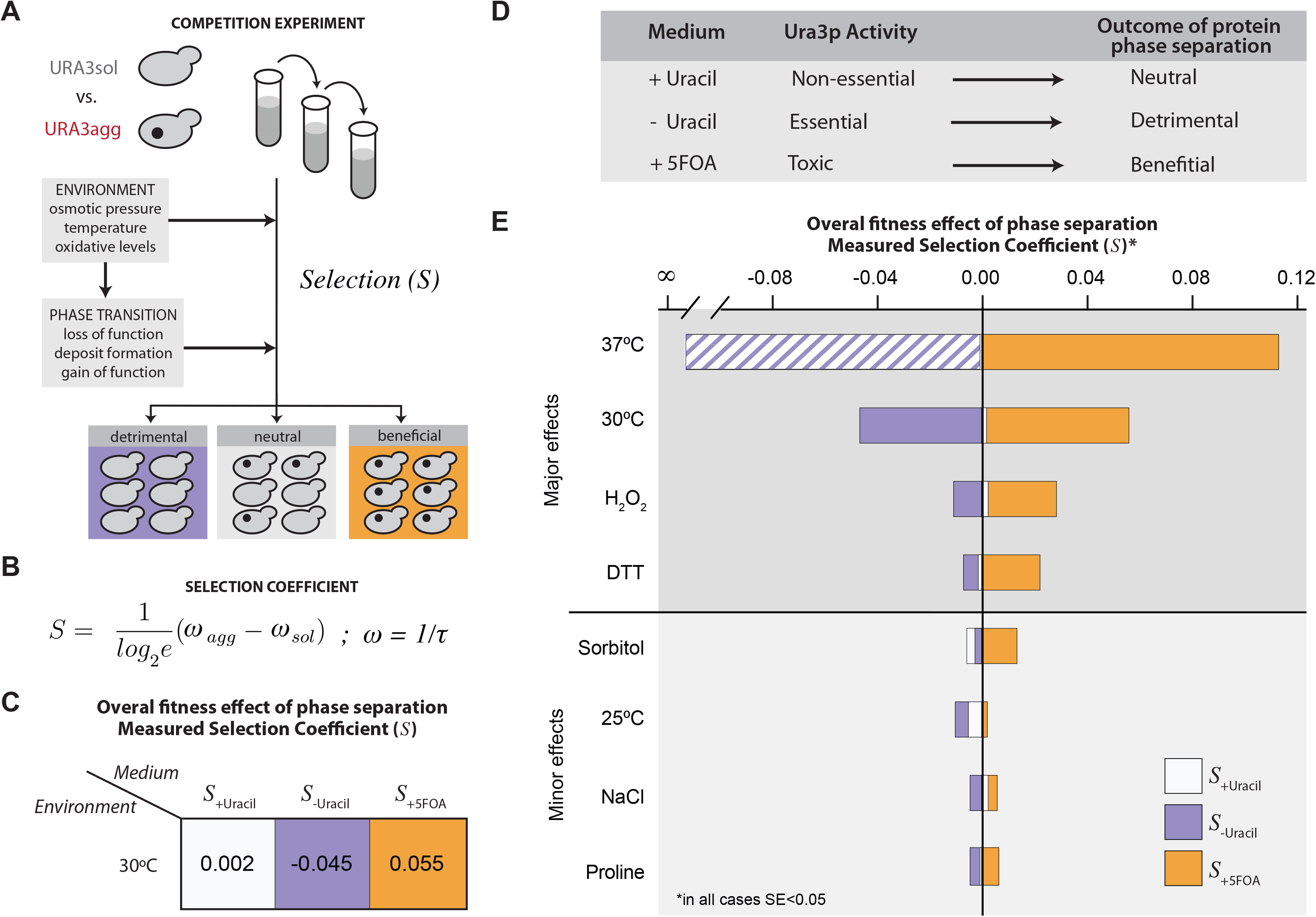
Protein phase separation and selection coefficient. **A.** Experimental design to measure the selection coefficient (*S*) upon growing URA3_agg_ in competition with URA3_sol_. **B.** The selection coefficient, *S*, is proportional to the difference in growth rates between URA3_agg_ and URA3_sol_ (**Material and Methods**). ω is the growth rate, which corresponds to the inverse of doubling time, τ. **C-D.** The *S* values measured at standard conditions (30°C) indicate that, depending on the media composition (+Ura, −Ura, +5FOA), the formation of protein deposits can be neutral, deleterious or even advantageous for the cell as the function becomes non-essential, essential or toxic, respectively. **E.** *S* values measured in different environments. In all cases the standard error is below 5%. See also **Appendix Table S1**. Color scale: purple in absence of uracil (-Ura), white in the presence of uracil (+Ura) and orange in the presence of 5FOA (+5FOA). At 37°C, the heat stress together with the inability to fold Ura3p have such a strong fitness effect that the strains barely grew in absence of uracil, impeding the measurement of *S*. A bar of purple-white diagonal lines indicates a presumed value of *S* at 37°C without uracil.

The results obtained at standard growth conditions (30°C) show that the formation of Ura3p_agg_ deposits can be neutral (in the presence of uracil; non-essential), deleterious (in the absence of uracil; essential) or even advantageous (in the presence of 5FOA; toxic) for yeast, depending on the composition of the growth medium (**Figure 2C**; average *S* values from two biological replicates). Therefore, without changing the protein sequence or the genotype we can quantify different overall effects of deposit formation on cell fitness (**Figure 2D**). In this manner, the system we developed allows for the quantification of the overall effects (both negative and positive fitness effects) of deposit formation upon protein phase separation in different environments (**Figure 2E**; **Appendix Table S1**). We find a wide range of selection coefficients suggesting that URA3_agg_ is differentially selected for or against when the role of the protein is different (essential/non-essential/toxic; in different media) and in different environments (**Figure 2E**). Changing the oxidation levels (0.5mM H_2_O_2_, 1mM DTT) or increasing the temperature (37°C) has a major effect on fitness compared to conditions in which we changed the osmotic pressure (1M Sorbitol, 0.5M NaCl), decreased the temperature (25°C) or when we added a chemical chaperone (0.5M Proline). This observation raises the question as to how and why does the phase separation of the same protein lead to such differences in selection coefficients under different environments? We investigate this question in the next sections (**Figure EV4**).

### Disentangling the different effects of protein phase separation

The overall fitness effect (cost/benefit) of protein phase separation is not only determined by the loss or gain of the biochemical activity of the protein, but also due to the cost of deposit formation (e.g. amino acid sequestration in deposits, sequestration of ATP dependent chaperone activity and other proteins, toxicity of the assembly, etc. (Gsponer & Babu, 2012; Maji et al, 2009; Olzscha et al; Patel et al, 2017; Sanchez de Groot et al, 2012; Suraweera et al, 2012; Tomala et al, 2014); **Figure 3A**, Eq.1). To infer the impact of these effects, we considered that the measured selection coefficient (*S*) is determined by a combination of three factors: (i) the cost of deposit formation, (ii) the cost of losing the essential Ura3p biochemical activity, and (iii) the benefit of gaining a protective function against the toxic activity by sequestering Ura3p in to deposits (**Figure 3A**, Eq.2). In conditions where Ura3p is not essential (with uracil), any change in cell fitness primarily depends on the fitness cost of deposit formation (**Figure 3A**, Eq.3). In the absence of uracil, the overall effect on fitness not only includes the cost of deposit formation but also the cost of reducing Ura3p activity due to deposit formation (**Figure 3A**, Eq.4). Finally, in the presence of 5FOA, the effect on fitness includes the cost of deposit formation, and the fitness benefit of reducing the toxic activity of Ura3p by sequestering the protein into the deposit (**Figure 3A**, Eq.5). Although some Ura3p activity can be present in deposits (O’Connell et al, 2014; Wallace et al, 2015), this is likely to be significantly reduced (e.g. due to protein conformational changes, and restricted access to substrate) compared to the free, diffusible well-folded protein as in the URA3_sol_ strain (Suresh et al, 2015).

**Figure 3.**
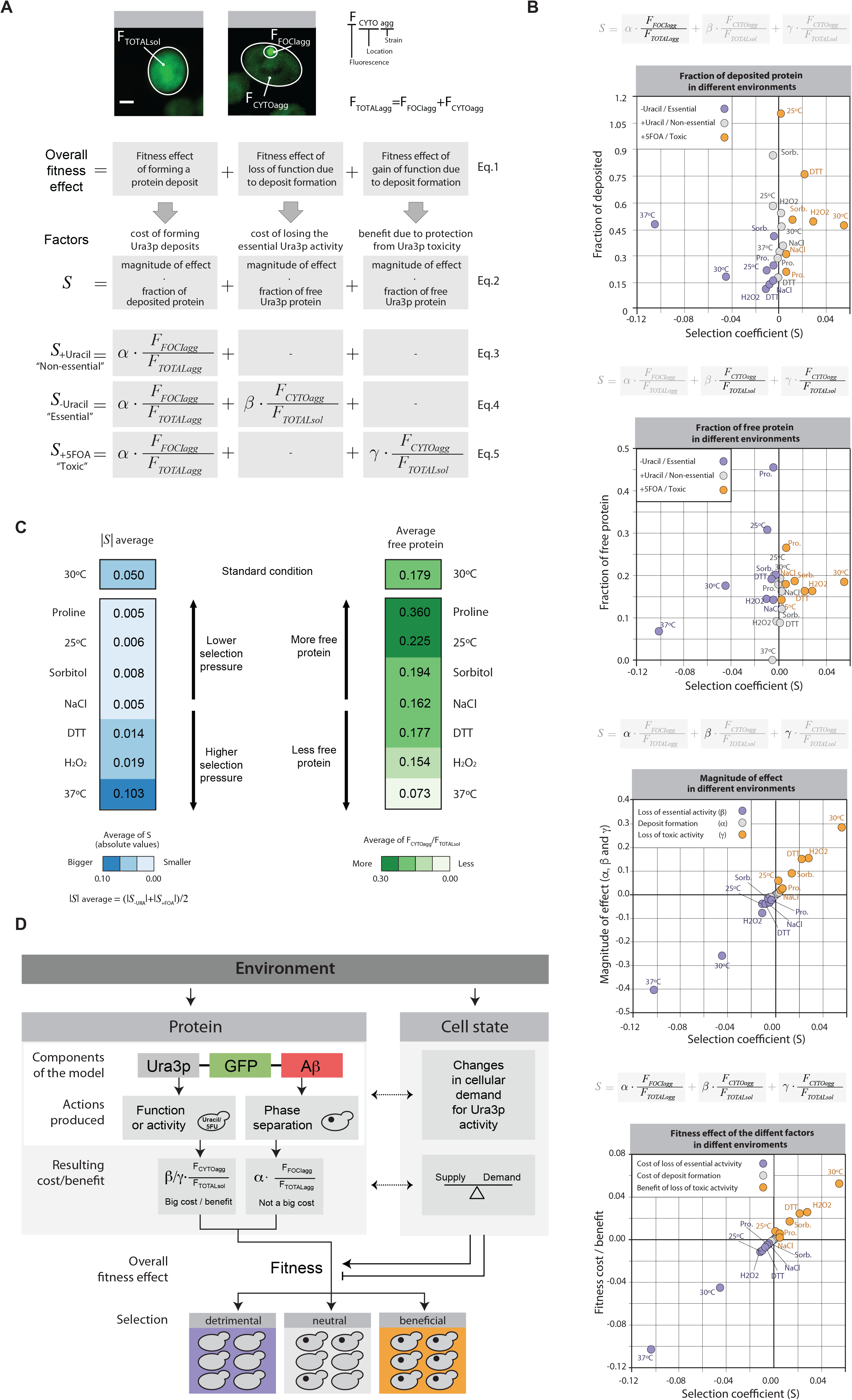
Fitness cost and benefit analysis. **A.** We measured the intensity of fluorescence emitted by the fluorescent reporter (GFP) to estimate: (i) the amount, (ii) the location and (iii) the deposited/diffused state of the model proteins. Accordingly, for each cell analyzed, we measured the fluorescence intensity of its cytoplasm (*F_TOTALsol_* for URA3_sol_ and *F_CYTOagg_* for URA3_agg_) and of its foci (*F_FOCIagg_*). We used these measurements to estimate the different effects of protein phase separation on cell fitness. The overall fitness effect of forming a protein deposit depends on three factors: foci formation, loss of function and gain of function (Eq.1). In our system, we can distinguish between the cost of forming Ura3p deposits, cost of losing the essential Ura3p activity and the benefit from protection against Ura3p toxicity. Each cost and benefit effect depend on the ratio of deposited or free protein and the specific environment. Accordingly, we can split these cost and benefit effects into two components: one associated with the amount of deposited or free protein (*F_FOCIagg_/F_TOTALagg_* or *F_CYTOagg_/F_TOTALsol_*) and a magnitude of effect defined by the environmental conditions (α, β or γ) (Eq.2). For the non-essential, essential and toxic roles (+Ura, −Ura and +5FOA) of Ura3p, the number and type of effects applicable is different (Eq.3, Eq.4 and Eq.5). **B.** Plots showing the distribution and relationship between the fractions of deposited or free Ura3p, its magnitude of effect on fitness (α/β/γ) and the selection coefficient (*S*) in different environments. **C.** Average of the selection coefficient (left) and average of the cytosolic fluorescence (equivalent to the fractional abundance of the free protein in URA3_agg_, right). **D.** Schematic representation highlighting how phase separation of the same protein can result in different fitness effects depending on the environment.

To relate the measured selection coefficient with the different effects of deposit formation, we quantified the total fluorescence (F_TOTALagg_) and the amount of fluorescence in deposit/foci (F_FOCIagg_), and the cytosol (F_CYTOagg_) for individual cells of the URA3_agg_ population using confocal microscopy (**Figure 3A**). The measured fluorescence is proportional to the amount of protein present in the cell (Soboleski et al, 2005). For the URA3_sol_ strain, the total fluorescence (F_TOTALsol_) is the same as cytosolic fluorescence (F_CYTOsol_) and is indicative of the maximum amount of free, active Ura3p that can be available in our designed system (**Material and Methods**). The cost of forming a deposit is defined as the product of the fraction of the protein in deposits (phase separated) (F_FOCIagg_/F_TOTALagg_) and a proportionality constant/coefficient (α) that depends on the environment (**Figure 3A** Eq.2 and Eq.3). This constant/coefficient is indicative of the magnitude of the effect of forming the protein deposit on the fitness in a particular environment, and is considered to remain the same for an environment irrespective of whether Ura3p is non-essential, essential or toxic (i.e. when grown in +Uracil, −Uracil or +5FOA; see **Material and Methods**). The effect of reduced Ura3p activity due to deposit formation is defined as the product of the apparent free, active protein (F_CYTOagg_/F_TOTALsol_, ratio of free protein in the URA3_agg_ strain to the maximum that can be available in a cell) and a proportionality constant/coefficient (β for cost / γ for benefit) that is indicative of the magnitude of the effect of the loss/gain of biochemical activity on fitness in a particular environment (**Figure 3A** Eq.4 and Eq.5, **Material and Methods**). Although both strains (URA3_agg_ and URA3_sol_) present similar transcript levels (**Figure EV2**), due to deposit formation, a fraction of translated Ura3p_agg_ can be rapidly removed through autophagy or other mechanisms (Sanchez de Groot et al, 2015; Villar-Pique & Ventura, 2013) (**Figure EV1**). This may hence lead to a reduction in F_TOTALagg_ when compared to F_TOTALsol_ (Sanchez de Groot et al, 2015; Villar-Pique & Ventura, 2013)(**Figure EV1**, **Appendix Figure S2**, **Appendix Table S1**, **Material and Methods**).

### Environment modulates the magnitude of the effects of phase separation on cell fitness

Environment can modulate the selection pressure on variants in a population; genotypes and characteristics that are beneficial in certain environments could become detrimental in others (Bershtein et al, 2012; Pena et al, 2010). We examined how the fraction of deposited vs. free Ura3p, the magnitude of effect, the fitness cost/benefit of deposit formation and the selection coefficient (*S*) varies in different environments.

When Ura3p activity is not essential for a cell, although the fraction of the deposited protein varies considerably in the different environments, the measured selection coefficient remains closer to 0 (0 means no difference in cell fitness, **Figure 3B**, first plot; grey dots). However, when Ura3p activity is essential or toxic, the fraction of the deposited protein in the different environments and the selection coefficient tend to vary considerably (e.g. *S* = −0.103, without uracil at 37°C and *S* = 0.055, with 5FOA at 30°C) (**Figure 3B**, first plot; purple and orange dots). Consistently, in terms of the fraction of free Ura3p, although the amount of free protein varies in the different environment, the measured selection coefficient does not differ much when Ura3p is not essential (**Figure 3B**, second plot; grey dots). When Ura3p is essential or toxic, the fraction of free protein in the different environments and the selection coefficient tend to vary considerably (**Figure 3B**, second plot; purple and orange dots). In addition, we find that the magnitude of effect for the cost of deposit formation (α) is considerably lower than the effects associated with protein function, both with loss of essential enzymatic activity (β) and gain of protection against the toxic activity (γ) in the different environments (**Figure 3B**, third plot). These observations quantify and reflect the essentiality and lethality associated with the biochemical reaction catalyzed by Ura3p. Moreover, whereas the cost of deposit formation remains relatively stable in the different environments (**Figure 3B**, fourth plot; grey dots), the fitness cost/benefit associated with reduced Ura3p activity varies significantly in different environments. Thus, in certain environments the amount of free Ura3p is more important for cell fitness than in others (e.g. 37°C and 25°C, without uracil, **Figure 3B**, second plot). In line with this observation, the environments with higher average absolute selection coefficient (**Figure 3C**, darker blue) tend to have lower average apparent free protein in the URA3_agg_ strain (**Figure 3C**, lighter green; Pearson Correlation Coefficient = −0.64).

Due to the modular nature of the designed protein, a change in the environment can: (i) affect the phase separation process (e.g. higher temperature can accelerate deposit formation (de Groot & Ventura, 2006) and/or (ii) affect the activity of Ura3p, for instance by enhancing protein folding (e.g. presence of the chemical chaperone proline (De Los Rios & Goloubinoff, 2012) or misfolding (e.g. higher levels of oxidative stress). Interestingly, environments that affect protein folding and hence Ura3p activity can directly influence the supply of nucleotides in a cell (**Figure 3D**). Moreover, the environment can also affect the cellular state, the growth rate and cause changes in the cellular demand for Ura3p activity (e.g. low temperature reduces speed of cell division, hence DNA replication and, thus, rate of nucleotide consumption; **Figure 3D**, right panel). In this context, it has been demonstrated that the growth rate affects cell fitness and the sensitivity to environmental stresses; for instance, rapidly growing cells tend to be more sensitive to antibiotics than slow growing or stationary cells (Berney et al, 2006; Lu et al, 2009). Thus, specific environments can influence protein folding, and hence the total amount of free/deposited protein, as well as modulate the cell state by influencing the supply/demand for a particular biochemical activity (**Figure EV4**). In this way, different environments can modulate the magnitude of the fitness costs and benefits of sequestering Ura3p_agg_ in deposits.

Overall, these results help explain why the phase separation of a single protein can lead to different selection coefficients in the different environments. Specifically, in our model protein, cell fitness primarily relies on the balance between the supply/demand of Ura3p activity and to a much lesser extent on the cost of deposit formation. This means that when uracil is present in the media, the cell does not require Ura3p (no demand). Hence a change in the amount (free/deposited) or quality of this enzyme (active, folded/inactive, misfolded) in a specific environment does not affect cell fitness significantly. In agreement with this possibility, in these environments, there is no correlation between the fraction of the deposited protein (or free protein) and the selection coefficient (**Figure 3B**, first and second panel, grey dots). However, when the protein activity is essential or toxic (i.e. in the absence of uracil or presence of 5FOA), each environmental condition has an associated Ura3p demand. Hence the amount of free protein (ratio of F_CYTOagg_/F_TOTALsol_, in other words the Ura3p supply) as well as the quality of the enzyme determines the overall fitness effect (Figure 3C-D).

Given the extensive variability in fitness of a strain expressing a protein that form deposits in different environments (**Figure 2** and **Figure 3**), we computationally investigated how the environment could influence the population frequency of variants carrying this protein at the population level.

### History of the environments determines population frequency of strains that can form phase-separated deposits

Our experiments were performed with a 1:1 population (URA3_agg_:URA3_sol_), grown in an exponential phase and with no resource limitation. However, in nature, variants that can form phase separated structures arise sporadically in an already growing population, possibly in more severe conditions (Newby & Lindquist, 2013). To investigate how such a variant changes in frequency with time in a population, and whether such a variant can take over the population in a defined time-scale, we developed a computational model and simulated a scenario where one individual in a population of a million cells acquires a mutation that result in the formation of protein deposits (**Figure 4A**; **Material and Methods**). Using the experimentally measured selection coefficient values (*S*) (**Appendix Table S1**), we followed how an individual that carries a protein prone to form deposits is selected for, or against in a cell population in diverse scenarios. From the simulations, we infer that in a stable environment and when deposit formation is beneficial, the variant can become a dominant member of the population (i.e. more than 50%) within ∼50-5000 generations (**Figure 4B** and **Figure 4C**, blue color range; **Appendix Table S2**). As expected, if protein phase separation is detrimental to cell fitness in an environment, the variants containing the protein that form deposits never achieve population frequencies higher than the starting condition, and get “diluted” with time (**Figure 4C**, orange color spectrum).

**Figure 4.**
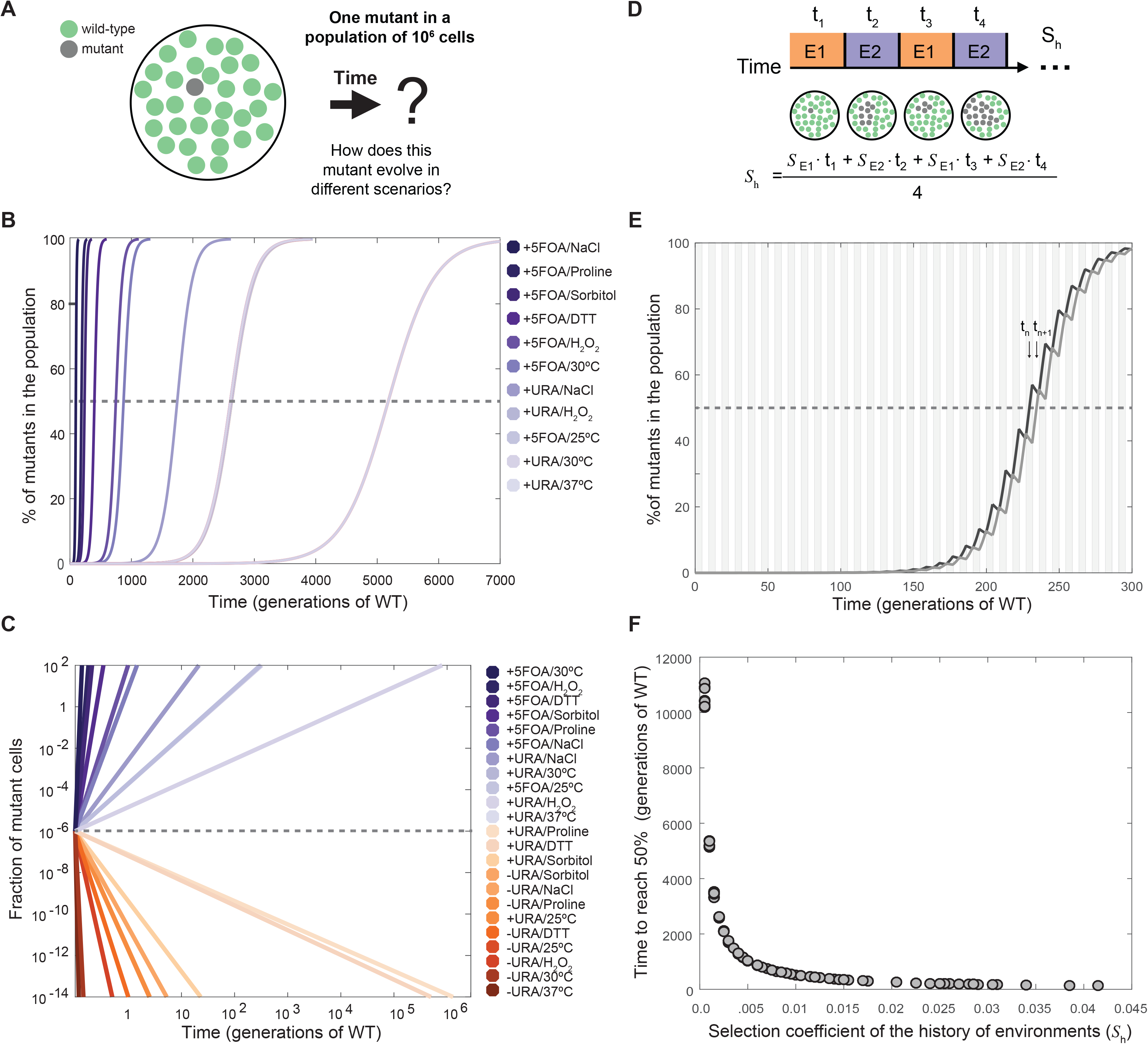
Computational simulation and estimation of population frequencies. **A.** Schematic of the question addressed for the computational simulation. **B.** Population frequency of the phase separation-prone variant over time (in number of generations with respect to wild-type strain at 30°C, 2.7 hours) in different non-oscillating conditions with a positive measured selection coefficient (*S*) for protein phase separation. A dashed line indicates the point where the mutant is present in 50% of the population. The corresponding value in the x-axis for each profile denotes the time required to reach this point. **C.** Population frequency of the phase separation-prone variant over time in different environmental conditions with positive and negative *S*. The dashed line indicates the initial fraction of mutant cells in the population and the colored lines indicated the fraction of mutant cells in the population at different time points (generations of WT is used as the unit of reference). **D.** Diagram showing how the frequency of the mutant cells can change in fluctuating environments (environment, E and time in environment, t). The sequence of environments (history) experienced by a mutant determines the overall selection coefficient (*S*_h_). **E.** Population frequency of the variant over time in two different alternating environments. Black line, profile of population frequency of the variant, where, at the moment in which the first mutant cell appears, the environment positively selects it; however, in the subsequent environment the selection is negative. Grey line, profile of population frequency of the variant, where, at the moment in which the first mutant cell appears, the environment negatively selects it; however, in the subsequent environment the selection is positive. The area of the plot is divided in grey (E1) and white (E2) sections to indicate the fluctuating environment with positive and negative *S*. **F.** History of environments resulting from any possible combination of two environments (e.g. “environment 1” 25°C and “environment 2” 0.5M Proline), from those experimentally tested in this work (actual data available in **Appendix Table S5**).

In nature, however, the growth environment can fluctuate more or less rapidly over time (e.g. day/night cycles affect growth temperatures daily) (Newby & Lindquist, 2013). This means that an individual with a protein variant that can phase separate can be selected to different extents as determined by the sequence/history of environments (*S*_h_) that the population experiences (Kussell & Leibler, 2005; Leibler & Kussell, 2010) (**Figure 4D**; **Appendix Table S3**). In rapidly fluctuating and alternating environments, as long as the average selection coefficient over the time (or selection coefficient of the history of environments, *S*_h_) is positive, the frequency of the phase separation-prone variant in the population becomes higher over time (**Figure 4E**). It is worth noting that the frequency of the variant becomes higher irrespective of whether the immediate environment when the mutation is acquired is beneficial (**Figure 4E**; dark line) or detrimental (**Figure 4E**; light line) to fitness, as long as the environment fluctuates and *S*_h_ is positive. Nevertheless, at a given time, the frequency of the phase separation-prone variant can be higher if the mutation is acquired in a favorable environment compared to a detrimental environment (**Figure 4E**; dark line vs. lighter line at t_n_ and t_n+1_). These simulations highlight that for the same population, the history of environments (*S*_h_) can determine the persistence and frequency of a phase separation-prone variant (**Figure 4F**).

### Protein phase separation: a source of phenotypic variability

Microscopy image analysis reveals the existence of stochastic, cell-to-cell variation in total protein abundance (i.e. expression noise (Acar et al, 2008; Burga et al, 2011; Chevin, 2011; Jothi et al, 2009; Ravarani et al, 2016; Soboleski et al, 2005) as measured by GFP fluorescence; **Figure 5A**, **Appendix Figure S3**). Under standard growth conditions (in which both strains present similar transcript levels, see **Figure EV2**), we find that the cell-to-cell variation in protein abundance is higher for URA3_agg_ compared to URA3_sol_ irrespective of whether Ura3p activity is essential, non-essential or toxic (as measured by the coefficient of variation; CV=100·σ/μ; ratio of the standard deviation to the mean abundance; **Figure 5B**). However, if we focus just on one genotype, regardless of URA3_sol_ or URA3_agg_, we obtain higher noise when uracil is added. This is consistent with the current view that the maximum population variance is achieved under conditions with no selection (Thompson & Cubillos, 2017). Interestingly, the cell-to-cell variation in protein abundance is higher in the deposits compared to the cytosol for URA3_agg_ irrespective of whether Ura3p is essential, non-essential or toxic (**Figure 5C**). These data suggest that the extent of protein deposit formation by phase separation can modulate the abundance of free/active protein in a cell (**Figure 5D**). We find that when Ura3p activity is essential or toxic, on average, cells with a higher fraction of phase-separated protein are selected against (i.e. higher free protein is selected for) or for (i.e. less free protein is selected for), respectively (**Figure 5D**). When Ura3p is not essential, the average foci size is in between what is observed for the other two growth media, suggesting no obvious selection for foci size (or free protein) (**Figure 5C**). These results indicate that even within an isogenic population, the extent of deposit formation between individuals can be selected for or against under different selective pressures. This effect is similar to what one observes in prion strain selection when yeast cells are grown under different stress conditions (Chakrabortee et al, 2016; Holmes et al, 2013).

In conditions where Ura3p activity is toxic, we observe that cells are elongated, tend to be larger and display cell division problems (**Figure 5E**). This is consistent with the fact that in the presence of 5FOA, free/active Ura3p produces 5FU and leads to cell cycle arrest during DNA synthesis (**Figure 5F**) (Seiple et al, 2006). Since the formation of phase separated structures reduces the amount of free/active protein (**Figure 5D**), individuals with larger amount of protein in deposits (F_FOCI_ > 70% F_TOTAL_) will have little free protein, and display less Ura3p activity (**Figure 5G**), and hence should less often exhibit cell cycle arrest. Similarly, individuals with smaller amount of protein in deposits will display higher Ura3p activity (**Figure 5G**), and hence should more often exhibit cell cycle arrest. In line with this expectation, we observe that within the same population, individual cells with more protein recruited into phase separated deposits tend to have less free protein after normalizing for cell area (**Figure 5H**). Furthermore, DNA quantification reveals that cells with more protein in phase separated deposits tend to display cell cycle arrest less often compared to those with less protein in deposits (**Figure 5I**). Indeed, in conditions where Ura3p activity is not essential no cell cycle arrest is observed (**Figure 5J**).

Taken together, our results suggest that the cell-to-cell variability in the extent of protein phase separation can lead to variability in the abundance of free, functional protein, which in turn can generate a phenotypic continuum in a genetically identical cell population (**Figure 6A**). It should be noted that depending on the molecular structure and the properties of the phase separated assembly (e.g. insoluble deposits or liquid droplets), one could find an opposite behavior where enzyme activity may be higher within certain types of phase separated structures (Shin & Brangwynne, 2017). Nevertheless, such phenotypic diversity, induced by variable protein phase separation, can provide a selective advantage under certain conditions and could enhance the likelihood of survival of some individuals from a population after an environment alteration. Similar benefits have been attributed to the prion conformational switch. An important difference, however, is that in the latter case, instead of generating a phenotypic continuum, the emergence of different prion strains provides new heritable phenotypes with distinct and largely non-overlapping phenotypes/functional states (Chakrabortee et al; Derdowski et al, 2010; Farkash-Amar et al, 2014; Halfmann et al, 2010; Holmes et al, 2013; Newby et al, 2017; Newby & Lindquist, 2013) (**Figure EV5**). Our observations suggest that, beyond prions, polypeptide chains with certain phase separation-/aggregation/deposit-forming propensity can contribute to selectable phenotypic heterogeneity to an isogenic cell population. In other words, the selectable phenotypic variability described for prions is likely to be a general property of any polypeptide segment that can form phase separated protein deposits.

**Figure 5.**
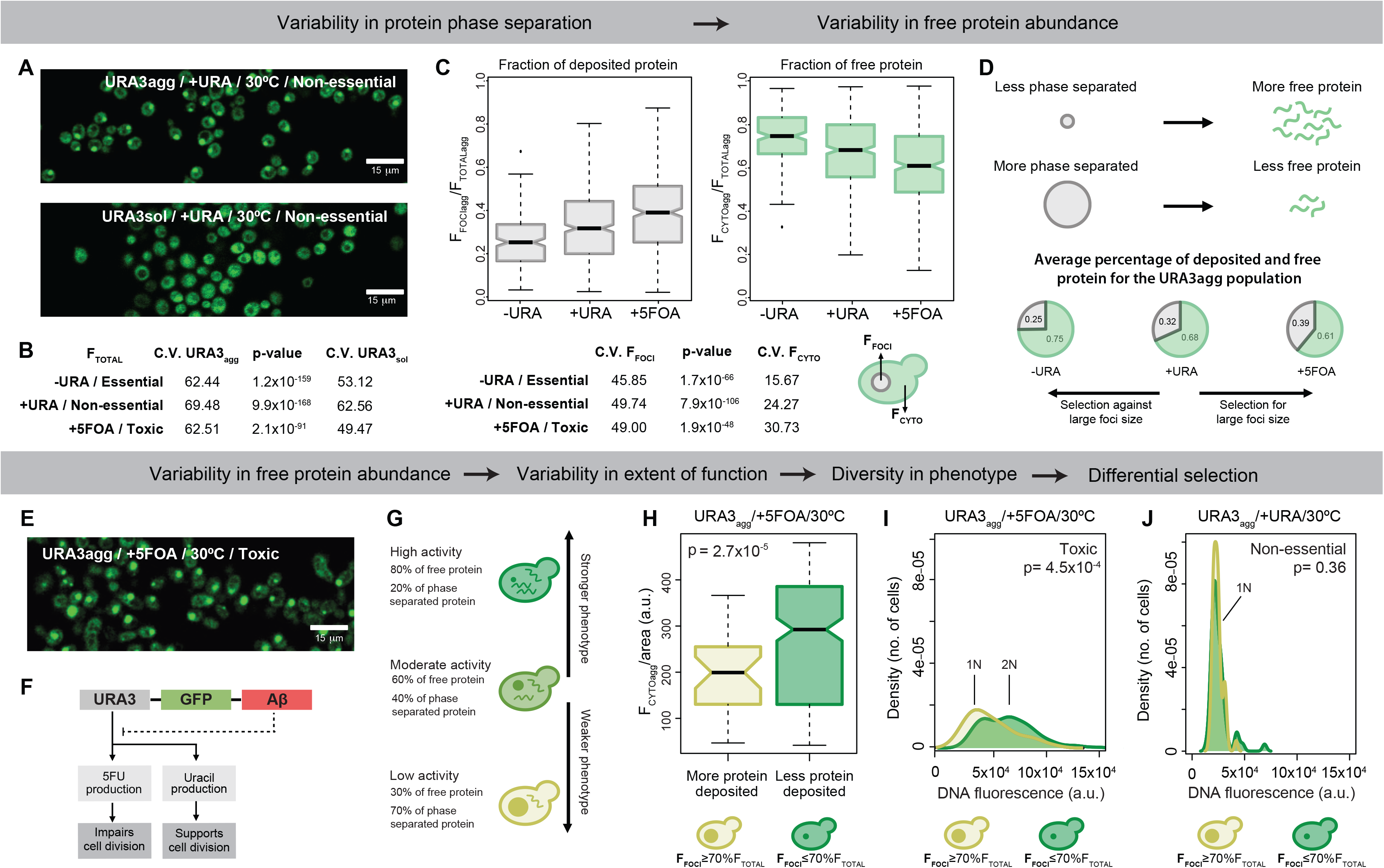
Impact on cell-to-cell variability and phenotypic diversity. Panels from **A** to **D** show that the variability in foci formation contributes to variability in free protein abundance. **A.** Confocal microscopy images of URA3_agg_ and URA3_sol_ incubated for 18 hours at 30°C. **B.** Coefficient of variation and p-values (Wilcoxon test) of the total fluorescence measured from populations of URA3_agg_ and URA3_sol_. **C.** Boxplot showing the distribution of foci fluorescence (left) or cytoplasmic fluorescence (right), from the foci containing subset of a population of URA3_agg_ grown at 30°C. Coefficient of Variation and p-values of each boxplot are shown below. **D.** Schematic showing how the formation of protein deposit can affect the abundance of free protein in a cell. Pie charts showing the average fraction of deposited and free protein of URA3_agg_ in different media composition and 30°C. **E.** Confocal microscopy image of URA3_agg_ incubated in the presence of 5FOA after 18 hours at 30°C. **F.** Schematic showing that in our system, the phase separation-promoting region can modulate Ura3p activity and thus its phenotypic effect in a cell. **G.** Schematic highlighting that higher/lower level of free protein inside the cell results in higher/lower Ura3p activity leading to the manifestation of a stronger/weaker phenotype. **H.** Boxplots of the cytoplasmic fluorescence normalized by the cell area (F_CYTOagg_/cell area) for individual cells of the URA3_agg_ population in 5FOA. Light green box, cells with foci containing at least 70% of the total fluorescence (big foci). Dark green box, cells with foci containing less than 70% of the total fluorescence (small foci). **I-J.** Cell cycle arrest phenotype. Density plots showing the fluorescence distribution of individual cells from the URA3_agg_ population stained with propidium iodide to measure their DNA content. The cells were incubated for 18 hours at 30°C without (**I**) and with 5FOA (**J**).

**Figure 6.**
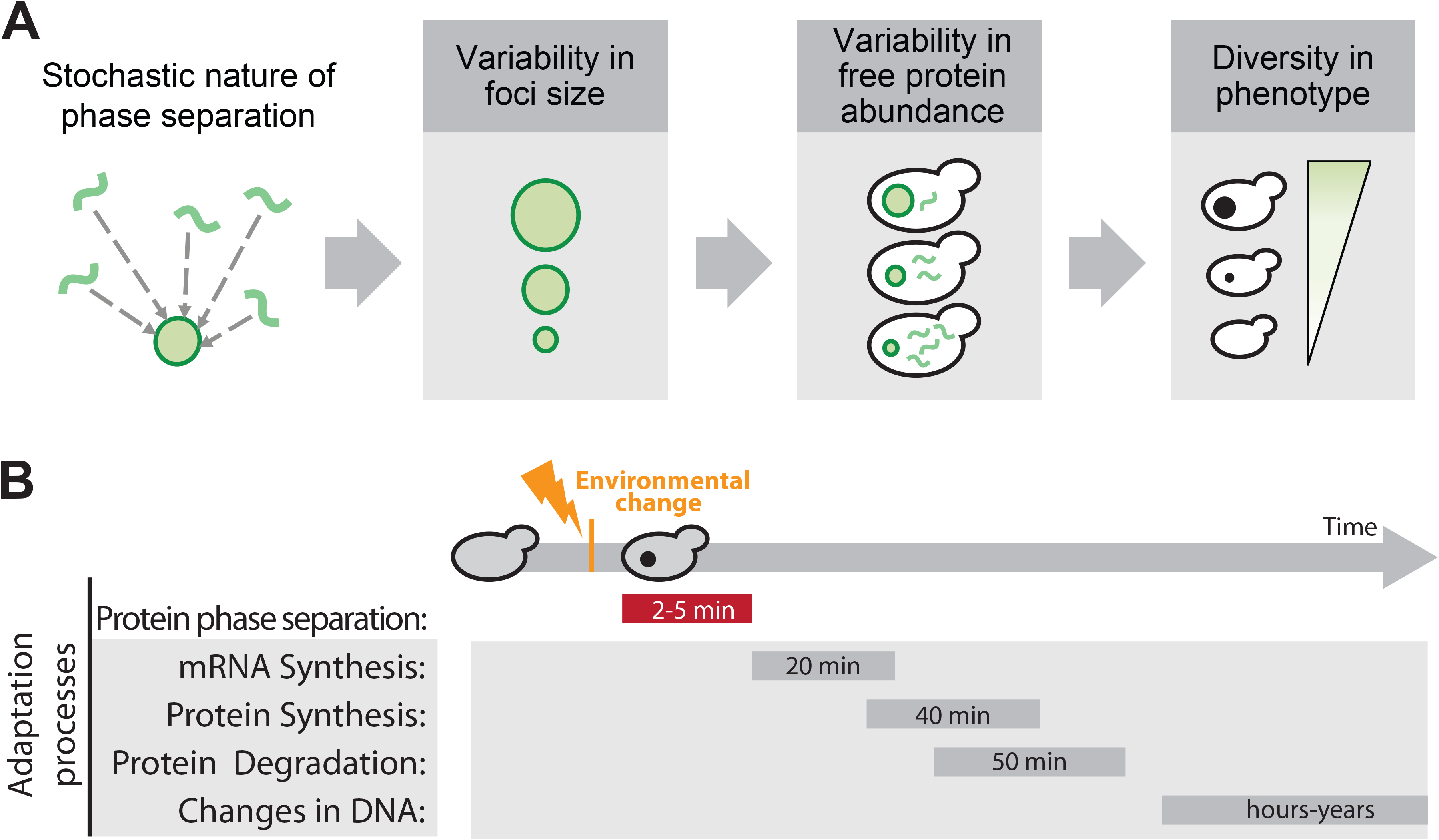
Phenotype modulators. **A.** The stochasticity of deposit formation due to phase separation will define the foci size, which in turn will influence the amount of free protein in the cytosol. The abundance of the free protein (or the phase separated structure) will ultimately determine the cellular phenotype. The behavior and phenotypic outcome may also be influenced by the molecular structure and properties of the phase separated structure (e.g. complex coacervates, liquid droplets, etc.) **B.** Timeline for the different processes that are important for adaptation/response to an environmental change. Time estimates were obtained from published datasets (Belle et al, 2006; Escusa-Toret et al, 2013; Wang et al, 2002).

## DISCUSSION

Our understanding of the process of protein phase separation has changed rapidly in the last decade. Currently, there is consensus that intracellular protein condensates are associated with a wide range of effects, ranging from pathological to beneficial (Aguzzi & Altmeyer, 2016). Clearly, not all deposits have the same molecular structure (Brangwynne et al, 2015; Feric et al, 2016; Lee et al, 2016; Molliex et al, 2015; Zhang et al, 2015). In the cellular milieu, the different types of deposits differ in their composition, dynamics and cellular location. This in combination with the biochemical function of the molecules that participates in phase separation determines the overall effect on fitness (Banani et al, 2016; Chavali et al, 2017b; Feric et al, 2016; Gsponer & Babu, 2012; Lin et al, 2015; Miller et al, 2015; Mitrea & Kriwacki, 2016; Molliex et al, 2015; Morley et al, 2002; Pak et al, 2016; Riback et al, 2017; Zhang et al, 2015; Zhu & Brangwynne, 2015). To investigate how phase separated protein deposits can affect fitness and to disentangle the effects of different factors on cell fitness, we studied a model protein with a modular architecture (**Figure 1B**) that is prevalent in nature (Bornberg-Bauer & Alba, 2013; Chavali et al, 2017a; Lees et al, 2016). Such a protein architecture may be widespread, possibly because it allows the properties of the polypeptide segment that promotes phase separation (e.g. concentration at which the protein forms deposits) to evolve independently of the biochemical function that is mediated by the rest of the protein. Proteins without this modular architecture may also tune their aggregation propensity; however, since function and aggregation propensity are linked in such proteins, evolution of their sequences will likely be more constrained.

Our work is consistent with previous studies (Escusa-Toret et al, 2013; Geiler-Samerotte et al, 2011; Tomala et al, 2014) and provides quantitative insights as well as a framework to decouple the fitness cost and benefit of protein phase separation, and the associated loss/gain of protein function in different environments. Overall, the data suggests that, in our system, yeast cells can tolerate and are adapted to intracellular protein phase separation of the model protein that we have designed. In future, this modular architecture can be adapted to assess the specific contribution of different types of phase separation promoting sequences on cell fitness. In our system, the effect on fitness is primarily driven either by the loss and/or gain of protein function as a consequence of sequestering the free/active/functional protein into insoluble deposits (**Figure 3C**). Furthermore, the environment can modulate the consequences of protein phase separation in strains with the same genomic background, enabling situations where protein deposit formation is either beneficial or detrimental to fitness (**Figure 2C**). These results indicate that the interplay between the protein state (e.g. soluble or deposited) and the functional demand placed by the cell defines the overall effect of protein phase separation on cell fitness. It also highlights that the specific growth environment is a key determinant of the effect of phase separation on cell fitness. Using computational simulations, we further showed that the frequency and persistence of a phase separation-prone variant within a population is determined by the history of environments experienced by this population (**Figure 4G**). In fact, it has been observed in nature that the history of environments (memory) can be ‘encoded’ by aggregation-prone Q/N-rich segments (Caudron & Barral, 2013; O’Connell et al, 2014; Wiltzius et al, 2009; Zhang et al, 2015).

Our observations on individual cells in an isogenic population indicate that the extent and onset of protein phase separation may vary between cells (**Figure 5**). While this may be influenced by different cellular processes (Chalancon et al, 2012) protein phase separation might further contribute to changes in free protein level and leads to cell-to-cell heterogeneity in free protein abundance. The existence of cytoplasmic chaperones that can dissolve protein macromolecular assemblies such as Hsp104p (Duennwald et al, 2012), and variability in their abundances between cells could further contribute to the ability of the phase separation-prone protein to be released or refold from deposits to become active. Such mechanisms that generate cell-to-cell variability in abundance can act in synergy and still be distinct from the ability of chaperones to help fold (i.e., buffer) deleterious mutants during or after protein synthesis (Burga et al, 2011), leading to stochastic changes in protein abundance between individuals.

A population should generate enough phenotypic variability to ensure that some cells will be able to survive an unexpected environmental change (Acar et al, 2008; Koonin, 2007; Kussell & Leibler, 2005; Leibler & Kussell, 2010). In this sense, like in the case of a prion conformational switch, phase separation can be a general mechanism of adaptation by resulting in rapid and transient manifestation of loss/gain of function compared to mechanisms that involve changes in the mRNA, or protein levels (Belle et al, 2006; Escusa-Toret et al, 2013; Kussell & Leibler, 2005; Leibler & Kussell, 2010; Wang et al, 2002) (**Figure 6B**). In this context, variation in the formation of deposits, and variable stoichiometry of the deposits between individuals of a population can further generate phenotypic diversity (**Figure 6A**). In other words, since deposit formation is dependent on protein concentration (Brangwynne et al, 2015; Ciryam et al, 2013; Veis, 2011), different individuals can form phase separated assemblies at different abundances by “sensing” their concentration in cells (Gsponer & Babu, 2012). This may lead to variable and rapid manifestation of phenotypes by transiently sequestering proteins into deposits (or restricting their localization in a cell) in different environments. Interestingly, in most cases this phenomenon is triggered because proteins have cellular concentrations above their saturation level (Ciryam et al, 2013). For these supersaturated proteins, formation of such deposits can be protective when toxic concentrations are reached in a cell (Bershtein et al, 2012).

The modular nature of the phase transition promoting-segment also helps with phenotypic diversity and the manifestation of new phenotypes during evolution, since such segments can evolve independently of the biochemical function mediated by the rest of the polypeptide (Boke et al, 2016; Bornberg-Bauer & Alba, 2013; Lees et al, 2016). This may possibly explain why many regulators such as transcription factors (e.g. Snf5) contain phase separation-promoting segments (e.g. Q/N rich segments) independently of structured domains that perform specific biochemical function such as DNA binding (Chavali et al, 2017a; Chong et al, 2018; Sabari et al, 2018). It is known that cell-to-cell variation in protein abundance of such regulators can lead to differential activation of downstream regulatory networks, and drive the phenotypic differences in an isogenic population (Gemayel et al, 2015; Halfmann et al, 2012; Holmes et al, 2013; Jothi et al, 2009). Thus, variability in deposit formation between individuals can lead to cell-to-cell differences in phenotypes by affecting the abundance of the free/active protein, as observed with foci size variability, and cell cycle arrest phenotype in the URA3_agg_ population (**Figure 5**).

The findings presented here suggests that in addition to variability in the process of gene expression, and protein folding between individuals of a population (Acar et al, 2008; Burga et al, 2011; Chevin, 2011; Holmes et al, 2013; Jothi et al, 2009; Kussell & Leibler, 2005; Leibler & Kussell, 2010; Ravarani et al, 2016), phase-separation induced cell-to-cell variability in free protein abundance is likely to be an important phenomenon that can facilitate adaptability in different conditions. Thus, studies on the effects of phase-separated structures such as intracellular protein deposition should not only involve a global investigation of the cost and benefit but also consider cell-to-cell variability of phenotypes in the population (Farkash-Amar et al, 2014).

Inside the cell, a myriad of proteins phase separate by different mechanisms. However, the cellular complexity does not end here since these assemblies can interact with each other, with other molecules and are subjected to a constant transformation due to the quality control machinery (Banani et al, 2016; Brangwynne et al, 2015; Farkash-Amar et al, 2014; Feric et al, 2016; Lin et al, 2015; Nott et al, 2015; Pak et al, 2016; Wu & Fuxreiter, 2016; Zhang et al, 2015; Zhu & Brangwynne, 2015). Overall, in an *in vivo* system, phase separation can affect the cellular processes at multiple levels and result in different fitness costs and benefits. In our system, the protein model phase separates from a largely soluble active state to a primarily insoluble inactive state. Obviously, in more complex systems (e.g. dynamic assemblies, active compartments, complex coacervates, etc.) there are more variables to be studied than those analyzed here. However, the quantitative insights disentangling the various effects, as well as the conceptual and methodological framework presented in this study may be adapted (e.g. by testing phase separation promoting sequences with different molecular structures and properties, adding new elements and in different genetic backgrounds) to understand the effect on cell fitness of phase separation of diverse proteins and how this determines their selection within a population.

## MATERIAL AND METHODS

### Strains and vectors

All the strains employed in this work are based on Y03157 (BY4741; Mat a; his3D1; leu2D0; met15D0; ura3D0; YBR020w::kanMX4) obtained from Euroscarf deletion collection. In this strain, the GAL1 gene is missing thereby hindering cells to consume Galactose (the inducer) as a carbon source (Geiler-Samerotte et al, 2011).

A multicloning site flanked by 65 bp region that is homologous to TRP1 (TRP start vector) was designed and introduced in a pMA vector (GeneArt® Life Technologies) (**Appendix**, Vectors and Primers). The TRP1 locus was selected as the insertion position for the proteins (Ura3p_sol_ and Ura3p_agg_), because of its close proximity to the centromere and the high expression levels of genes in this region. These properties minimize the frequency of gene silencing and ensure a steady state expression level through several generations. A yeast-optimized GFP codon and the SpHis5 selection marker were extracted from a pKT128 vector (Sheff & Thorn, 2004) and inserted between the restriction sites PacI and SacI. The pGAL1 promoter was obtained from a pESC-URA vector (Agilent Technologies) and inserted between HindIII and SalI. The same vector was used to extract the locus encoding the URA3 gene plus an additional 12 residues linker (Linker 1: GGTACCGCTAGTGGTTCTGCTGGTTCTGCGATTAAC). URA3 was inserted between pGAL1 and GFP using the In-Fusion® HD Cloning Kit (Clontech) (TRP-URA vector).

The sequence of Aβ42 was optimized for yeast using the codon usage reported in the Codon Usage Database (http://www.kazusa.or.jp/codon/) and was built using four DNA oligos. Aβ42 was inserted 3’prime of the GFP (TRP-URA-AB vector) using the In-Fusion® HD Cloning Kit (Clontech) and included a 12 residues linker in between (Linker 2: GGTGGAAGTGCTAATGGTACTTCTGGTGCTAGTGGT).

Both linkers were designed to be yeast codon optimized and enriched in small amino acids to provide flexibility and avoid steric hindrances. Additionally, these sequences have a low net charge and no aggregation prone regions to prevent interference with the phase separation process triggered by Aβ42. This second property was measured using the AGGRESCAN algorithm (Conchillo-Sole et al, 2007; de Groot et al, 2012). The cellular behavior of Aβ42 has been extensively studied in bacteria and yeast, both by others and us (de Groot et al, 2006; de Groot & Ventura, 2006; Morell et al, 2011; Sanchez de Groot et al, 2015; Villar-Pique et al, 2012). In yeast, the formation of deposits/granules appears to protect cells against the toxic species (e.g. soluble oligomers) (de Groot & Ventura, 2006; Sanchez de Groot et al, 2015; Villar-Pique & Ventura, 2013).

### Rationale and design of model proteins

In this work, we designed model proteins to study the effects of protein phase separation on cell fitness. With this aim we designed a polypeptide with a modular arrangement, similar to those recently found in nature (Kato et al, 2012; Kedersha et al, 2013; McGlinchey et al, 2011; Sleeman & Trinkle-Mulcahy, 2014) (**Figure 1C**). This setup allows us to tune different properties while studying the cost/benefit of the phase separation event (see Main Text).

To build our model proteins, we chose the yeast endogenous Ura3p enzyme as the functional domain. This provides a versatile framework for assessing the different roles of the enzyme, as the biochemical activity can be essential, non-essential, or toxic depending on the composition of the growth medium.

To report the expression, distribution and location of the fusion protein we added the enhanced GFP (F64L/S65T) as the reporter. This GFP variant includes a mutation at the position 64 (F−>L) that allows its correct production at 37 °C (Cormack et al, 1996; Day & Davidson, 2009). This extra stability is crucial for the present work to ensure the integrity of the reporter under stress conditions. Importantly, as reported in earlier studies in *E. coli*, even under strong overexpression, the folding of this enhanced GFP fused with phase separation-promoting peptide is faster than the phase separation process itself. This property results in the formation of fluorescent foci at a temperature range between 25 and 37°C (de Groot et al, 2006; de Groot & Ventura, 2006). Furthermore, to distinguish between the effect of the environment and foci formation on the steady state abundance of the model protein (GFP fluorescence), our system includes a soluble control (URA3_sol_) to indicate the maximum amount of active GFP that can be made available in a cell at a certain environment. Hence, in the unlikely event in which a GFP molecule will misfold due to reasons not associated with the phase separation process, the normalization against the control strain will account for such a situation.

Finally, Aβ42 was added as a tag to promote phase separation and study the effect of deposit formation. This peptide is the main component of the plaques found in patients with Alzheimer’s disease (Chiti & Dobson, 2006; Ciryam et al, 2013; Olzscha et al, 2011; Pak et al, 2016; Woerner et al, 2016). *In vitro*, Aβ42 assembles into amyloids fibrils driven by its hydrophobic residues. When overexpressed in unicellular models this peptide accumulates into inclusion bodies (de Groot et al, 2006; de Groot & Ventura, 2006; Escusa-Toret et al, 2013; Morell et al, 2011; Pak et al, 2016; Plata et al, 2010; Sanchez de Groot et al, 2015; Villar-Pique et al, 2012; Villar-Pique & Ventura, 2013). Aβ42 has no intrinsic function in *S. cerevisiae* and, according to our previous studies, its expression has almost no effect on yeast growth, at least under optimal grow conditions (Sanchez de Groot et al, 2015; Villar-Pique & Ventura, 2013). Consistent with this, it is worth emphasizing that we measured no significant difference in growth that was associated with the addition of Aβ42 in conditions where URA3p is non-essential and no stress is introduced (**Figure 2**). An alternative strategy to induce the phase separation of Ura3p could be the alteration of its sequence with the aim to destabilize the protein and to induce foci formation. However, this mutant variant can also affect Ura3p structure and enzymatic activity, increasing the complexity of the system and making it more difficult to determine the origin of the cell fitness alterations. We also considered the addition of a less aggregation prone Aβ42 peptide, however all point mutations studied have a similar growth rate (Sanchez de Groot et al, 2015; Villar-Pique & Ventura, 2013) and to obtain a fully soluble variant, multiple mutations are necessary (Wurth et al, 2002). Therefore, to generate the soluble version of the model protein, we used the version without a tag. In short, we designed the model protein in a modular manner with the addition of a phase separation promoting tag (i.e. separate functional and aggregation prone region in the same polypeptide).

With the above-mentioned elements, we built two different constructs: (i) one was designed to control for the intrinsic fitness cost of overexpressing a soluble protein (Geiler-Samerotte et al, 2011; Kafri et al, 2016; Plata et al, 2010) (Ura3p fused to GFP, Ura3p_sol_) and (ii) the other one was designed to investigate the effect of protein deposit formation (Ura3p fused to GFP and Aβ42 peptide, Ura3pagg, **Figure 1D**). By comparing the overall cell fitness upon expressing these two constructs, we obtain the fitness effects of cells expression the phase separated Ura3p_agg_. It is worth mentioning that the obtained phenotype is not that of a full null strain. The microscopy images demonstrate that there is still cytosolic fluorescence in the Ura3p_agg_ strain (**Figure 5** and **Figure EV1**). These data suggest that the fitness effects associated with deposit formation of Ura3p are smaller than those obtained with a true auxotroph (Mulleder et al, 2012).

For the mathematical modelling of this system (see selection coefficient calculation) we have considered some generalizations for the analysis of the different parameters influencing the phase separation event and its effect on cell fitness. We have subtracted the protein exchange between the phase-separated assembly and the surrounding milieu from our analyses based on two experimental observations. First, FRAP analyses show that Ura3p_agg_ forms a stable, non-dynamic deposit similar to IPODs (**Figure EV3**). Second, our analyses are done keeping the culture at exponential phase and after long expression times (from 18 hours to 3 days). This set up allows Aβ42-GFP deposits to reach an equilibrium in which most cells contain just one big focus (see **Figure 5**) and the amount of protein synthesized/degraded appears to be constant (Morell et al, 2011). Thus in these conditions we consider that the cells reach a “population equilibrium” in which some properties of the culture are constant: (**a**) the number of new born cells is constant (exponential phase); (**b**) the cells are adapted to the environment (after 18-72 hours) and therefore the environmental pressure/selection is constant; (**c**) and considering assumptions **a** and **b** the noise distribution of Ura3p expression, and the number of foci forming cells in the population remains stable (Morell et al, 2011; Sanchez de Groot et al, 2015). This premise also considers that the cell-to-cell heterogeneity in foci formation is a final outcome of the inherent variability of cellular processes such as protein expression and quality control activity (e.g. chaperones) (Chalancon et al, 2012; Farkash-Amar et al, 2014; Ravarani et al, 2016).

Under the microscope Ura3p_sol_ seems homogeneously distributed throughout the cytosol whereas part of Ura3p_agg_ is accumulated in to foci. It has been reported that not all the protein in the soluble fraction is active (e.g. soluble oligomers) and that not all the aggregate protein is inactive (i.e. fluorescence into foci) (Villar-Pique & Ventura, 2013). However, at least in our system, due to the static nature of the analyzed foci, the Ura3p recruited in them will have limited access to its substrate (irrespective of its conformation and functional state). In addition, theoretically, the free Ura3p should have less constraints to acquire a correct fold than the protein located in the foci (Morell et al, 2011). Overall, based on these premises, we assume that the protein located in the cytosolic fraction is more active and accessible to substrate than the protein located in the assembly; so, we consider that the residual activity located in the foci is negligible. It should be noted that depending on the nature of the phase separated structure (e.g. liquid droplets), one could find an opposite behavior where enzyme activity may be higher within certain types of phase separated structures (Shin & Brangwynne, 2017).

### Media composition and environmental conditions

In all the assays, the strains were grown in SD -HIS media containing a mixture of sugars and amino acids. Since protein concentration is a critical factor for foci formation, to guarantee high levels of expression we measured the effect of Galactose concentration (the inducer) on URA3_sol_ expression by monitoring the GFP fluorescence using flow cytometry (**Appendix Figure S2D**). Maximum levels of expression were obtained between 0.5 and 2%. Below 0.5% of Galactose, the fluorescence declines abruptly, so we chose 1% as a reliable concentration to keep high levels of induction. To obtain a fresh culture before inducing Ura3p expression, single colonies were picked and grown overnight in 2% Glucose. This culture was employed to inoculate SD -HIS 2% Raffinose and growth during six hours. Then it was inoculated in fresh media with 2% Raffinose and 1% Galactose. SD -His -Ura was employed to test the essentiality of Ura3p. SD -His with uracil and 5FOA (Zymo Research) was used to analyze the phase separation effects of a toxic Ura3p activity.

In all cases, the media was adjusted to pH 4.5 to ensure (when required) the 5FOA permeability. To test different environments, the different compounds were added before the pH adjustment. The final concentrations tested are: 0.5 mM H_2_O_2_, 1.5 mM DTT, 0.5 M NaCl, 1 M Sorbitol and 0.5 M Proline. The cells were grown at 220 rpm and 30°C before starting the experiments to test the effect of a specific environment Subsequently the cultures were incubated at 30° with the corresponding concentration of the above-mentioned chemical or without the chemical but shifting the temperature to 25° or 37°C.

### Cell cycle arrest

Cells grown until exponential phase were fixed by incubating them during 1 hour with ethanol 70% at 4°C. Then, the cells were treated with 1 mg/ml RNase A (Sigma) for 2 hour at

37 °C. The samples were stained with propidium iodide (50 µ g/ml) for 1 hour before analyzing them under a confocal microscope. The fluorescence of propidium iodide was employed as a measure of DNA content and an indication of the cell cycle arrest.

### Confocal microscopy

Cells were grown at exponential phase with 1% Galactose and the corresponding environmental condition for 18-20 hours before acquiring the images. The images were acquired with a Zeiss710 (Carl Zeiss) with an objective of 63x, an excitation laser of 488 nm and emission window between 581 nm and 750 nm. The cells expressing soluble Ura3p (Ura3p_sol_) display fluorescence that is homogeneously distributed through the cytoplasm whereas the cells that additionally express the Aβ (Ura3pagg) showed fluorescence in foci. At least 100 cells were captured for each strain and environment. For the propidium iodide assay, the fluorescence was excited with 561 nm and the emission was collected between 566 nm and 719 nm.

### Image processing

All the images were processed with Fiji (ImageJ). For each cell the program measured the cell area, the cytosolic fluorescence and the fluorescence contained in the aggregates. A macro was created to perform these measures automatically. The parameters of this macro for a whole cell analyses the presence of elements with: i) a fluorescent intensity between 20 and 255 a.u., ii) an area (size) between 8 and 50 μm^2^ and iii) a circularity between 0.25-1. To detect foci, a mask was generated including the elements with a fluorescent intensity between m2 40 and 255 a.u. and an area between 0.2 and 10 μm^2^. For the cells expressing Ura3pagg, the fluorescence from the foci was subtracted from the total fluorescence to obtain the amount of soluble protein in each cell. The cell size analysis shows a positive correlation with the foci size (**Appendix Figure S3**), probably due to aging and yeast asymmetric division, which retains proteins deposits in mother cells (that tend to bigger) producing clean daughter cells (that tend to be smaller) where there protein assembly has to start *de novo* (Coelho et al, 2014). For the propidium iodide assay, we adapted the macro to additionally measure the fluorescence in the propidium iodide channel for each element detected.

### Immunochemistry

Yeast cells were grown for 18 h in media with 2% Raffinose and 1% Galactose. 20 ml of culture was divided in two and centrifuged. For total fraction a pellet was resuspended in 75ml of lysis buffer (10 mM Tris-HCl pH 1/4 8, 150 mM NaCl, 0.05% Tween 20, 10% glycerol, 5 mM EDTA, 1 mM DTT, 2 mM PMSF) and 25 ml NuPAGE LDS sample buffer with 2.5% 2-mercaptoethanol (w/w) and incubated at 100°C for 5 min. For fraction separation a pellet was resuspended in 75 ml Y-Per yeast protein extraction reagent (Thermo Scientific) supplemented with 0.1mM PMSF and incubated at room temperature with agitation for 20 min. Then, the sample was centrifuged to separate the two fractions: soluble (supernatant) and insoluble (pellet resuspended in 75 ml of PBS). 25 ml of NuPAGE LDS sample buffer with 2.5% 2-mercaptoethanol (w/w) was added to both fractions and then incubated at 100°C. To separate the proteins, 5 ml of the total fraction and 10 ml of soluble or insoluble fractions were eluted into a precast NOVEX NuPAGE 4-12% gels in denaturing conditions. The Invitrogen iBlot system was used to transfer proteins to PVDF membranes. After blocking, membranes were incubated overnight at 4°C with anti-GFP rabbit antibody (Santa Cruz sc-8334) or anti-PGK1 mouse antibody (Novex 459250) diluted 1:1,000 and 1:10,000 respectively. Secondary incubation with anti-Protein G HRP conjugated (Millipore 18-161) at 1:10,000 was performed at RT during one hour. Images were taken with a Amersham Imager 600. ImageJ 2.0.0-rc-48 software was used to quantify protein bands.

### In vivo half-life measurement

Yeast cells were grown for 18 h in media with 2% Raffinose and 1% Galactose. Protein production was then stopped by cleaning the cells and changing the media to SD -HIS with 2% Glucose. All samples were adjusted to the same concentration before monitoring the turbidity (absorbance, 600 nm) and fluorescence (excitation 480 nm, emission 510 nm) with a TECAN infinite 200. Samples were agitated during 10 seconds before each measurement and during 450 seconds between time points. The fluorescence was measured as the ratio between fluorescence and turbidity at every time point. To calculate the ratio of fluorescence loss, the data were fitted to a Boltzmann’s sigmoid with GraphPad PRISM 5 software (GraphPad Software).

### Competition experiments

Cells pre-induced with 1% Galactose were grown in SD -His overnight and then inoculated into fresh media for 4 hours to reach exponential phase. Then the OD_600nm_ of the cultures was measured and the two strains were mixed in equal concentration (1:1) and this mixture was employed to inoculate the different media. At this point, we took a sample for time 0. At least two different duplicates of each culture were analyzed. The cells were kept at exponential phase to guarantee a stable duplication time and to minimize the number of old and dead cells. For each culture, samples were collected at six different time points (0h, 17h, 34h, 51h, 58h, 75h, 82h). At each time point the cells were centrifuged and the genomic DNA was extracted. After taking the samples, a dilution was performed to maintain the cultures at OD_600nm_ between 0.005 and 0.6.

To quantify the proportion of each strain, a pair of oligos (solF/solR and aggF/aggR) was designed to obtain a specific product close to 100 bp (**Appendix**, Vectors and Primers). The oligos to quantify URA3_sol_ anneal between URA3 and GFP. For URA3_agg_, the primers anneal specifically to the Aβ sequence. The specificity and efficiency of each pair of oligos was tested against the plasmids containing the inserted cassettes (**Figure EV2**). A control PCR with an empty plasmid (TRP start) produced no amplification product. For each qPCR assay, two standard slopes of plasmids encoding for Ura3p_sol_ and Ura3p_agg_ (pESC-URA and TRP-URA-AB vectors) were included in the same plate to correct for PCR efficiency and measure the number of gene copies (**Figure EV2**). The ΔΔCT method was employed to analyze each qPCR reaction (Teste et al, 2009). Log_2_ of the ratio of the strains (URA3_agg_/URA3_sol_) was plotted. The resultant slope corresponds to the difference between the growth rates of the two strains.

Before the optimization of the above-described method of growth measurement we also tried to implement other approaches previously reported as highly efficient to measure the effects of protein misfolding on cell fitness (Geiler-Samerotte et al, 2011). However, several experimental problems appeared after altering the environmental conditions. For example, we were not able to follow by flow cytometry the competition between yeast strains expressing a soluble and insoluble protein variant tagged with different fluorescent markers (i.e. YFP and GFP) at 37°C or 0.5 mM H_2_O_2_. The extreme growth conditions employed influenced the marker’s folding and fluorescence spectrum, impeding the correct identification of the populations.

### Doubling time

URA3_sol_ and URA3_agg_ were grown separately in 96 well plates. The turbidity (OD_600nm_) and fluorescence (450 nm excitation & 510 nm emission) were recorded at 72 hours with a Tecan Infinite M200 Pro. Before starting the assay, the cells were pre-induced with SD -His 2% Raffinose 1% Galactose overnight and then incubated in fresh media for 4 hours to reach exponential phase. The time course assay was started with an OD_600nm_ of 0.02 (Cary50 Bio). A spline interpolation approach was applied with the RStudio program to measure the slope of each growth curve, resulting in the doubling time τ (**Appendix Table S2**).

### Selection coefficient calculation

The selection coefficient is defined as in Chevin (Chevin, 2011):

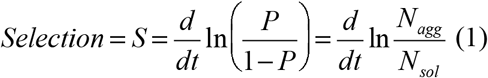

where P is the proportion of a certain strain in the population, N_agg_ is the number of cells of URA3_agg_ and N_sol_ is the number of cells of URA3_sol_ at time *t*.

The proportion of each strain at a certain time point is:

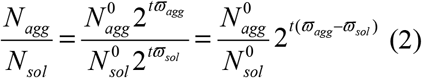

where N^0^ is the initial concentration (concentration at time 0) and ω is the growth rate. The measurement could be simplified using the base two logarithm and adding the difference between the growth ratio of the two strains, which can be obtained from the competition experiments:

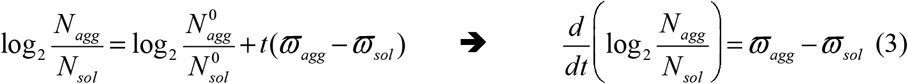

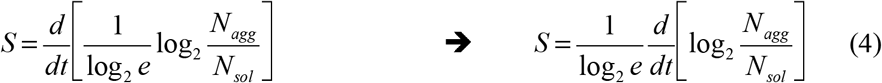

By combining (3) and (4), the selection coefficient could be calculated as:

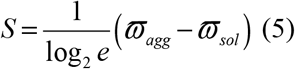

### Population frequency simulation

The doubling times (τ) employed in the simulations were interpolated from the selection coefficient values (*S*) and the experimentally measured doubling times of URA3_sol_ (the wild-type) (see **Appendix Table S1** and **Appendix Table S2**).

To simulate the time evolution of the mutant fraction in a mixed population, we started with a mixed population consisting of one mutant cell (URA3_agg_) and 10^6^ wild-type cells (URA3_sol_). In the simulation, we used the logarithmic scale to handle large numbers, but the final numbers were converted to linear scale whenever needed. We used an Euler integration scheme (with Δt = 60 min) to numerically calculate number of cells, N(t), at any given time, t:

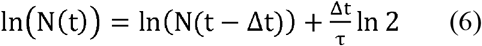

where τ is the doubling time (τ=1/ϖ). To simulate alternating environments, we used the same integration scheme as above, but the doubling times were re-assigned to match a specific environment, i.e. every 12 hours. All simulations were performed in MATLAB.

### Measurement of the effect of protein phase separation

In our experiments, the effects of Ura3p phase separation on cell fitness is determined by three factors: (i) the cost of deposit formation, (ii) the cost of losing the essential Ura3p biochemical activity, and (iii) the benefit of gaining a protective function by sequestering Ura3p in the deposit. The effect of foci formation comprises of the cost due to the sequestration of material and consumption of energy (e.g. amino acid sequestration, ATP dependent chaperone activity, etc.). The other two factors are associated with protein function that existed before foci formation (loss of function) and the generation of new beneficial effect after foci formation (gain of protection against the toxic function). In our experiments, the loss of function leads to a decrease in pyrimidines (in the absence of uracil), whereas the gain of protective function is due to the prevention of toxic 5-fluorouracil production (in the presence of 5FOA). We consider that each effect is determined by the amount of phase-separated protein (F_FOCIagg_/F_TOTALagg_), the amount of soluble protein (F_CYTOagg_/F_TOTALsol_) and a coefficient of proportionality that captures the effect of the specific environment (α, β, γ) (**Figure 3**, see main text). These coefficients indicate the magnitude of the effect on fitness of forming the deposit, loss and gain of protein function due to phase separation in a particular environment.

For a particular environment, we consider α to remain the same irrespective of whether Ura3p is non-essential, essential or toxic (i.e. when grown in +Uracil, -Uracil or +5FOA). In conditions where Ura3p activity is neither essential nor toxic we consider that there is no loss or gain of function and that the differences between URA3_agg_ and URA3_sol_ are a consequence of the phase separation process. Thus, the cost of protein deposit formation due to phase transition could be expressed as:

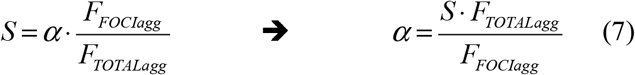

Where F_FOCIagg_ is the fluorescence contained in the deposits and F_CYTOagg_ is the florescence from the cytoplasm. The ratio quantifies the extent of phase separation. α is the magnitude of the effect of forming the deposit in specific environment.

By considering that α remains the same for a specific environment when grown in the different growth media, we can estimate the magnitude of effect for the loss (β) and gain (γ) of function when grown in the absence of uracil, and in the presence of 5FOA, respectively. Therefore, when Ura3 activity is essential the magnitude of effect of the loss of protein function due to phase separation can be expressed as:

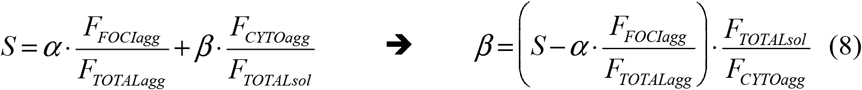

Where F_TOTALsol_ is the fluorescence of the cells expressing Ura3p_sol_ measured in the corresponding environment. F_TOTALsol_ represents the 100% of active enzyme that is available to a cell in a particular environment for a non-aggregation prone Ura3p_sol_. F_CYTOagg_/F_TOTALsol_ measures the amount of activity that is lost in the aggregating strain.

In conditions where Ura3 activity is toxic (i.e. uracil and 5FOA is added in the medium), the magnitude of effect of the gain of protection against the toxic activity (γ) could be calculated as below:

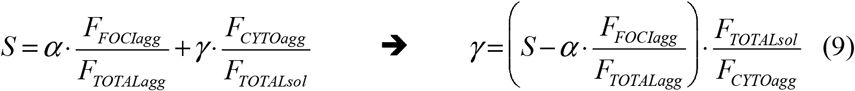

The three effects can be integrated in one general equation that could be applied for any of the environments and Ura3p role (essential, non-essential, toxic) analysed (Figure 3):

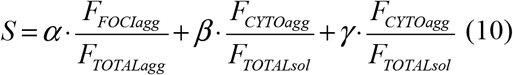

### mRNA expression levels

5 ml of fresh cells grown in 1% Galactose were centrifuged and suspended in 1 ml of 0.2 M Lithium Acetate 1% SDS solution. After 5 minutes at 70°C, 3 ml of TRIzol® was added. Next, the RNA was extracted with TRIzol® Reagent following the instructions provided by Life Technologies and the concentration of purified mRNA was measured. Retrotranscription was performed using the RevertAid H Minus First Strand cDNA Synthesis Kit (Thermo scientific) and the random hexamer primers included with the kit. The concentration of the cDNA generated was adjusted before running the qPCR experiment. The reactions were done using the SYBR® Green PCR Master Mix (Life technologies) and a Viia7 (AppliedBiosytems). The pair of primers (FmRNA/RmRNA) was designed using primerBLAST to anneal to the GFP region and not to the yeast genome (**Appendix**, Vectors and Primers). The mRNA of a reference gene (ALG9) was measured to normalize the data. The primers to amplify this gene were obtained from Teste *et al*.(Teste et al, 2009). Transcript quantification was done using the ΔCt method where the variation between URA3sol and URA3agg was calculated as ΔΔCt. The measured fold change between URA3sol and URA3agg is 1.07.

## DATA SOURCES

We provide all relevant datasets at the Appendix section.

## DATA AVAILABILITY

We provide all relevant datasets at the Appendix section.

**Expanded View** for this article is available online.

## ACKNOWLEDGEMENTS

This work was supported by the Medical Research Council (MC_U105185859; M.M.B., M.T., C.R., N.S.G.), the Center of Models for Life through Danish National Research Foundation (A.T.), Marie Curie Actions (FP7-PEOPLE-2012-IEF-330352; M.T. and FP7-PEOPLE-2011-IEF299105; N.S.G), FEBS Long-Term Fellowships (N.S.G.), Beatriu de Pinos fellowships (M.T.), the Ministerio de Economía y Competitividad (SAF2017-82158-R; SAF2015-72518-EXP and RYC-2012-09999; M.T.). N.S.G. was a recipient of the MRC centenary award. M.M.B. is a Lister Institute Research Prize Fellow. We thank Pavithra Chavali, Xiaohan Li, Greet De Baets, Natasha Latysheva, Sreenivas Chavali, Guilhem Chalancon, Louis Maddox and Anthony Fitzpatrick for critically reading this paper. We thank Gian Gaetano for supporting N.S.G and Sean Munro for the gift of strain Y03157.

## AUTHOR CONTRIBUTIONS

M.M.B., N.S.G. and S.V. conceived the project. M.M.B. and N.S.G. designed the project. M.T. and C.R. helped with experimental design. N.S.G., M.T. performed the experiments. A.T. performed the computational simulations. N.S.G. and M.T. analyzed the data and performed the computational and statistical analysis. M.M.B. and N.S.G. wrote the manuscript and all authors were involved in discussion and preparation of the final manuscript. N.S.G. led the project and M.M.B. supervised the project.

## EXPANDED VIEW FIGUREURE LEGENDS

**Figure EV1. Protein levels and half-life of URA3_sol_ and URA3_agg_**. **A.** Analysis of the expression and the soluble/insoluble part of URA3_sol_ and URA3_agg_. Total protein fraction obtained by two independent experiments after 18 hours of induction. The number indicates the URAp concentration (upper bands) quantification normalized by the concentration of PGK1p (lower bands). URA3_sol_ distribution is 88% soluble and 12% insoluble. URA3_sol_ is 6% soluble and 94% insoluble. Band analysis have been done with ImageJ. **B.** *In vivo* degradation rate was measured as the fluorescence loss after induction inhibition (see Material and Methods section). The plot shows the fluorescence normalized by the concentration of cells (absorbance at 600 nm) at each time point. The slope (absolute value), standard error and r-square of three technical replicates obtained from two different experiments are shown. The data was fitted to a Boltzmann’s sigmoid.

**Figure EV2. Variation in transcript expression and PCR efficiency. A.** URA3sol and URA3agg have similar mRNA abundance according to the ΔΔCt method (Teste et al, 2009). These measurements were made with a pair of primers that anneal to a common region located in the GFP sequence, named primers FmRNA and RmRNA respectively. The standard curves obtained for these primers and for **B** those used for the population quantification (solF/solR primers for URA3sol cells and the aggF/aggR primers for URA3agg) are also shown (**Material and Methods**).

**Figure EV3. Properties of URA3_agg_ deposits using FRAP.** We tested the consistency of the URA3_agg_ foci by photobleaching one half of the deposit (green line and green square) and monitoring the protein diffusion from the other half (blue line and blue square) with confocal microscopy. The assay was monitored for 40 seconds and shows no fluorescence loss or gain in any of the sides. This indicates that these foci are very dense, very much like an insoluble protein deposit (IPOD) (Kaganovich et al, 2008). **A.** Foci before (Time 0s) and after (Time 40s) photo-bleaching. **B.** Graph showing the fluorescence changes along time in each side of the foci. **C-D.** We also tested the conformation of the protein enclosed in the URA3_agg_ foci by analyzing the binding of anti-oligomer antibodies (Rabbit Anti-Oligomer (A11) AHB0052). As a secondary antibody, we used Goat Anti-Rabbit IgG H&L tagged with an Alexa Fluor® 555 (ab150078). We obtained a bright fluorescence signal that colocalizes with the foci’s GFP fluorescence, indicating that the deposits are rich in oligomeric structures. As a control, we also incubated URA3_sol_ with these antibodies obtaining few faint Alexa Fluor foci at the cytoplasm, indicating the absence or very low presence of oligomers.

**Figure EV4. Summary of the effects associated with URA3_agg_ phase separation in different environments that affect the topology of Ura3p, and hence protein activity (supply) as well as the cell state (demand). A**. Diagram showing different activity associated with the free and deposited protein. **B**. Summary table of the effects associated with URA3p_agg_ deposition and their effects on the selection outcome when growing in competition with URA3_sol_.

**Figure EV5. Population diversity introduced by prion conformational switching and by deposit formation.** Prion conformational switching is associated to the emergence of different prion strains that provide new heritable phenotypes with distinct and largely non-overlapping phenotypes or functional states. Whereas, a more general protein deposit formation upon phase separation may create a phenotypic continuum.

