## Supplemental Figures and Tables for "The fitness cost and benefit of phase separated protein deposits"

**Appendix**

**Table of contents**

**Vectors and Primers**	2

**Appendix Table S1**. List of data measured for each strain, growth media and environment.	6

**Appendix Table S2**. List of doubling times measured for each strain, growth media and environment.	7

**Appendix Table S3**. Time required to reach 50% of the population in (simulated) alternating environments.	8	11

**Appendix Figure S1**. Domain organization of proteins that can undergo phase separation and form functional deposits in vivo

**Appendix Figure S2**. Variation in fluorescence depending on the strain and the concentration of inducer.	13

**Appendix Figure S3**. Cell area vs. foci size distribution for URA3_agg_ cells grown in different environments when Ura3p is non-essential (+URA).	17

**Appendix References**	19

**Vectors and Primers**

TRP start sequence

(The regions homologous to TRP1 are indicated in bold)

5’GGCGCGCCGCGGCCGCGATATC**GAGGGCATTGGTGACTATTGAGCACGTGAGTATACGTGATTAAGCACACAAAGGCAGCTTGGAGT**CTCGAGGCGAGACGTCATCGAAGCTTACCGACTAGTACGAGAGCTCAGCCATGCATGCGAGAATTC**AAATGGCGTTATTGGTGTTGATGTAAGCGGAGGTGTGGAGACAAATGGTGTAAAAGACTCTAACA**GCTAGCGCCGGCGGTACC 3’

Restriction map of TRP start

```
                                                                                                             SacII
                                             BssHII NotI
                                            AscI     EagI   EcoRV
    CGAATTGGCGGAAGGCCGTCAAGGCCACGTGTCTTGTCCAGGCGCGCCGCGGCCGCGATA
  1 ---------+---------+---------+---------+---------+---------+
    GCTTAACCGCCTTCCGGCAGTTCCGGTGCACAGAACAGGTCCGCGCGGCGCCGGCGCTAT

                                   AccI
    TCGAGGGCATTGGTGACTATTGAGCACGTGAGTATACGTGATTAAGCACACAAAGGCAGC
 61 ---------+---------+---------+---------+---------+---------+
    AGCTCCCGTAACCACTGATAACTCGTGCACTCATATGCACTAATTCGTGTGTTTCCGTCG
                   BsmBI                                       SphI
           XhoI      AatII     HindIII   SpeI      SacI      NsiI
    TTGGAGTCTCGAGGCGAGACGTCATCGAAGCTTACCGACTAGTACGAGAGCTCAGCCATG
121 ---------+---------+---------+---------+---------+---------+
    AACCTCAGAGCTCCGCTCTGCAGTAGCTTCGAATGGCTGATCATGCTCTCGAGTCGGTAC

           EcoRI
    CATGCGAGAATTCAAATGGCGTTATTGGTGTTGATGTAAGCGGAGGTGTGGAGACAAATG
181 ---------+---------+---------+---------+---------+---------+
    GTACGCTCTTAAGTTTACCGCAATAACCACAACTACATTCGCCTCCACACCTCTGTTTAC

                      NheI  NaeI  KpnI  PacI
    GTGTAAAAGACTCTAACAGCTAGCGCCGGCGGTACCTTAATTAATGGAGCACAAGACTGG
241 ---------+---------+---------+---------+---------+---------+
    CACATTTTCTGAGATTGTCGATCGCGGCCGCCATGGAATTAATTACCTCGTGTTCTGACC


    CCTCATGGGCCTTCCGCTCACTGC
301 ---------+---------+----
    GGAGTACCCGGAAGGCGAGTGACG
```

Fragment of TRP-URA

5' muticloning site and barcode:	1 - 130

GAL1 promoter:	131 - 595 (blue)

URA3 gene:	602 - 1402 (green)

Linker 1:	1403 - 1438

GFP gene:	1439 - 2155 (violet)

TEF terminator:	2163 - 2392 (red)

SpHis5 marker cassette:	2782 - 3668 (orange)

3' muticloning site and barcode:	3669 - 3813

5'GGCGCGCCGCGGCCGCGATATCGAGGGCATTGGTGACTATTGAGCACGTGAGTATACGTGATTAAGCACACAAAGGCAGCTTGGAGTCTCGAGCGTTGATGCTCTACAAGATGCCGTGCTGCAGAAGCTT**TGAAGTACGGATTAGAAGCCGCCGAGCGGGTGACAGCCCTCCGAAGGAAGACTCTCCTCCGTGCGTCCTCGTCTTCACCGGTCGCGTTCCTGAAACGCAGATGTGCCTCGCGCCGCACTGCTCCGAACAATAAAGATTCTACAATACTAGCTTTTATGGTTATGAAGAGGAAAAATTGGCAGTAACCTGGCCCCACAAACCTTCAAATGAACGAATCAAATTAACAACCATAGGATGATAATGCGATTAGTTTTTTAGCCTTATTTCTGGGGTAATTAATCAGCGAAGCGATGATTTTTGATCTATTAACAGATATATAAATGCAAAAACTGCATAACCACTTTAACTAATACTTTCAACATTTTCGGTTTGTATTACTTCTTATTCAAATGTAATAAAAGTATCAACAAAAAATTGTTAATATACCTCTATACTTTAACGTCAAGGAGAAAAAACCCCGGATCC**GTCGAC**ATGTCGAAAGCTACATATAAGGAACGTGCTGCTACTCATCCTAGTCCTGTTGCTGCCAAGCTATTTAATATCATGCACGAAAAGCAAACAAACTTGTGTGCTTCATTGGATGTTCGTACCACCAAGGAATTACTGGAGTTAGTTGAAGCATTAGGTCCCAAAATTTGTTTACTAAAAACACATGTGGATATCTTGACTGATTTTTCCATGGAGGGCACAGTTAAGCCGCTAAAGGCATTATCCGCCAAGTACAATTTTTTACTCTTCGAAGACAGAAAATTTGCTGACATTGGTAATACAGTCAAATTGCAGTACTCTGCGGGTGTATACAGAATAGCAGAATGGGCAGACATTACGAATGCACACGGTGTGGTGGGCCCAGGTATTGTTAGCGGTTTGAAGCAGGCGGCAGAAGAAGTAACAAAGGAACCTAGAGGCCTTTTGATGTTAGCAGAATTGTCATGCAAGGGCTCCCTATCTACTGGAGAATATACTAAGGGTACTGTTGACATTGCGAAGAGCGACAAAGATTTTGTTATCGGCTTTATTGCTCAAAGAGACATGGGTGGAAGAGATGAAGGTTACGATTGGTTGATTATGACACCCGGTGTGGGTTTAGATGACAAGGGAGACGCATTGGGTCAACAGTATAGAACCGTGGATGATGTGGTCTCTACAGGATCTGACATTATTATTGTTGGAAGAGGACTATTTGCAAAGGGAAGGGATGCTAAGGTAGAGGGTGAACGTTACAGAAAAGCAGGCTGGGAAGCATATTTGAGAAGATGCGGCCAGCAAAAC**GGTACCGCTAGTGGTTCTGCTGGTTCTGCGATTAAC**ATGTCTAAAGGTGAAGAATTATTCACTGGTGTTGTCCCAATTTTGGTTGAATTAGATGGTGATGTTAATGGTCACAAATTTTCTGTCTCCGGTGAAGGTGAAGGTGATGCTACTTACGGTAAATTGACCTTAAAATTTATTTGTACTACTGGTAAATTGCCAGTTCCATGGCCAACCTTAGTCACTACTTTAACTTATGGTGTTCAATGTTTTTCTAGATACCCAGATCATATGAAACAACATGACTTTTTCAAGTCTGCCATGCCAGAAGGTTATGTTCAAGAAAGAACTATTTTTTTCAAAGATGACGGTAACTACAAGACCAGAGCTGAAGTCAAGTTTGAAGGTGATACCTTAGTTAATAGAATCGAATTAAAAGGTATTGATTTTAAAGAAGATGGTAACATTTTAGGTCACAAATTGGAATACAACTATAACTCTCACAATGTTTACATCATGGCTGACAAACAAAAGAATGGTATCAAAGTTAACTTCAAAATTAGACACAACATTGAAGATGGTTCTGTTCAATTAGCTGACCATTATCAACAAAATACTCCAATTGGTGATGGTCCAGTCTTGTTACCAGACAACCATTACTTATCCACTCAATCTGCCTTATCCAAAGATCCAAACGAAAAGAGAGACCACATGGTCTTGTTAGAATTTGTTACTGCTGCTGGTATTACCCATGGTATTGATGAATTGTACAAATAA**GGCGCGCC**ACTTCTAAATAAGCGAATTTCTTATGATTTATGATTTTTATTATTAAATAAGTTATAAAAAAAATAAGTGTATACAAATTTTAAAGTGACTCTTAGGTTTTAAAACGAAAATTCTTATTCTTGAGTAACTCTTTCCTGTAGGTCAGGTTGCTTTCTCAGGTATAGTATGAGGTCGCTCTTATTGACCACACCTCTACCGGCAGATCCGCTAGGGATAACAGGGTAATAT**AGATCTGTTTAGCTTGCCTCGTCCCCGCCGGGTCACCCGGCCAGC**GACATGGAGGCCCAGAATACCCTCCTTGACAGTCTTGACGTGCGCAGCTCAGGGGCATGATGTGACTGTCGCCCGTACATTTAGCCCATACATCCCCATGTATAATCATTTGCATCCATACATTTTGATGGCCGCACGGCGCGAAGCAAAAATTACGGCTCCTCGCTGCAGACCTGCGAGCAGGGAAACGCTCCCCTCACAGACGCGTTGAATTGTCCCCACGCCGCGCCCCTGTAGAGAAATATAAAAGGTTAGGATTTGCCACTGAGGTTCTTCTTTCATATACTTCCTTTTAAAATCTTGCTAGGATACAGTTCTCACATCACATCCGAACATAAACAACCATGGGTAGGAGGGCTTTTGTAGAAAGAAATACGAACGAAACGAAAATCAGCGTTGCCATCGCTTTGGACAAAGCTCCCTTACCTGAAGAGTCGAATTTTATTGATGAACTTATAACTTCCAAGCATACAAACCAAAAGGGAGAACAAGTAATCCAAGTAGACACGGGAATTGGATTCTTGGATCACATGTATCATGCACTGGCTAAACATGCAGGCTGGAGCTTACGACTTTACTCAAGAGGTGATTTAATCATCGATGATCATCACACTGCAGAAGATACTGCTATTGCACTTGGTATTGCATTCAAGCAGGCTATGGAGTAACTTTGCCGGCGTTAAAAGATTTGGACATGCTTATTGTCCACTTGACGAAGCTCTTTCTAGAAGCGTAGTTGACTTGTCGGGACGGCCCTATGCTGTTATCGATTTGGGATTAAAGCGTGAAAAGGTTGGGGAATTGTCCTGTGAAATGATCCCTCACTTACTATATTCCTTTTCGGTAGCAGCTGGAATTACTTTGCATGTTACCTGCTTATATGGTAGTAATGACCATCATCGTGCTGAAAGCGCTTTTAAATCTCTGGCTGTTGCCATGCGCGCGGCTACTAGTCTTACTGGAAGTTCTGAAGTCCCAAGCACGAAGGGAGTGTTGTAAAGAGTACTGACAATAAAAAGATTCTTGTTTTCAAGAACTTGTCATTTGTATAGTTTTTTTATATTGTAGTTGTTCTATTTTAATCAAATGTTAGCGTGATTTATATTTTTTTTCGCCTCGACATCATCTGCCCAGATGCGAAGTTAAGTGCGCAGAAAGTAATATCATGCGTCAATCGTATGTGAATGCTGGTCGCTATACTGCTGTCGATTCGATACTAACGCCGCCATCC**AGTTTAAACGAGCTCCATGTGGTTGCCTTACGCCGACTACGGATCCATGCATGCGAGAATTCAAATGGCGTTATTGGTGTTGATGTAAGCGGAGGTGTGGAGACAAATGGTGTAAAAGACTCTAACAGCTAGCGCCGGCGGTACC 3'

Fragment of pMA vector (residues 3814 – 6183 from TRP-URA vector):

5'ttaattaatggagcacaagactggcctcatgggccttccgctcactgcccgctttccagtcgggaaacctgtcgtgccagctgcattaacatggtcatagctgtttccttgcgtattgggcgctctccgcttcctcgctcactgactcgctgcgctcggtcgttcgggtaaagcctggggtgcctaatgagcaaaaggccagcaaaaggccaggaaccgtaaaaaggccgcgttgctggcgtttttccataggctccgcccccctgacgagcatcacaaaaatcgacgctcaagtcagaggtggcgaaacccgacaggactataaagataccaggcgtttccccctggaagctccctcgtgcgctctcctgttccgaccctgccgcttaccggatacctgtccgcctttctcccttcgggaagcgtggcgctttctcatagctcacgctgtaggtatctcagttcggtgtaggtcgttcgctccaagctgggctgtgtgcacgaaccccccgttcagcccgaccgctgcgccttatccggtaactatcgtcttgagtccaacccggtaagacacgacttatcgccactggcagcagccactggtaacaggattagcagagcgaggtatgtaggcggtgctacagagttcttgaagtggtggcctaactacggctacactagaagaacagtatttggtatctgcgctctgctgaagccagttaccttcggaaaaagagttggtagctcttgatccggcaaacaaaccaccgctggtagcggtggtttttttgtttgcaagcagcagattacgcgcagaaaaaaaggatctcaagaagatcctttgatcttttctacggggtctgacgctcagtggaacgaaaactcacgttaagggattttggtcatgagattatcaaaaaggatcttcacctagatccttttaaattaaaaatgaagttttaaatcaatctaaagtatatatgagtaaacttggtctgacagttaccaatgcttaatcagtgaggcacctatctcagcgatctgtctatttcgttcatccatagttgcctgactccccgtcgtgtagataactacgatacgggagggcttaccatctggccccagtgctgcaatgataccgcgagaaccacgctcaccggctccagatttatcagcaataaaccagccagccggaagggccgagcgcagaagtggtcctgcaactttatccgcctccatccagtctattaattgttgccgggaagctagagtaagtagttcgccagttaatagtttgcgcaacgttgttgccattgctacaggcatcgtggtgtcacgctcgtcgtttggtatggcttcattcagctccggttcccaacgatcaaggcgagttacatgatcccccatgttgtgcaaaaaagcggttagctccttcggtcctccgatcgttgtcagaagtaagttggccgcagtgttatcactcatggttatggcagcactgcataattctcttactgtcatgccatccgtaagatgcttttctgtgactggtgagtactcaaccaagtcattctgagaatagtgtatgcggcgaccgagttgctcttgcccggcgtcaatacgggataataccgcgccacatagcagaactttaaaagtgctcatcattggaaaacgttcttcggggcgaaaactctcaaggatcttaccgctgttgagatccagttcgatgtaacccactcgtgcacccaactgatcttcagcatcttttactttcaccagcgtttctgggtgagcaaaaacaggaaggcaaaatgccgcaaaaaagggaataagggcgacacggaaatgttgaatactcatactcttcctttttcaatattattgaagcatttatcagggttattgtctcatgagcggatacatatttgaatgtatttagaaaaataaacaaataggggttccgcgcacatttccccgaaaagtgccacctaaattgtaagcgttaatattttgttaaaattcgcgttaaatttttgttaaatcagctcattttttaaccaataggccgaaatcggcaaaatcccttataaatcaaaagaatagaccgagatagggttgagtggccgctacagggcgctcccattcgccattcaggctgcgcaactgttgggaagggcgtttcggtgcgggcctcttcgctattacgccagctggcgaaagggggatgtgctgcaaggcgattaagttgggtaacgccagggttttcccagtcacgacgttgtaaaacgacggccagtgagcgcgacgtaatacgactcactatagggcgaattggcggaaggccgtcaaggccacgtgtcttgtcca 3'

Fragment of TRP-URA-AB

Linker of 12 amino acids (GGSANGTSGASG) (Supplemental Experimental Procedures) and fragment of Aβ was added between GFP and the TEF terminator. Insertion site after residue 2152 of TRP-URA vector (uppercase Linker 2 and bold Aβ42 sequence):

5'GGTGGAAGTGCTAATGGTACTTCTGGTGCTAGTGGT**gatgctgaatttagacatgattctggttatgaagttcatcatcaaaaattggtcttttttgctgaagatgttggttctaataaaggtgctattattggtttgatggttggtggtgttgttattgct** 3'

Sequences of the primers used

Primers for qPCR, to monitor the competition assay:

solF: TCCAAACGAAAAGAGAGACCACA

solR: AGTGGCGCGCCTTATTTGTA

aggF: TGGTGCTAGTGGTGATGCTG

aggR: AGAACCAACATCTTCAGCAAAAA

Primers to measure the production of transcripts:

FmRNA: TAGTCACTACTTTAACTTATGGTGTTCAA

RmRNA: CTTTCTTGAACATAACCTTCTGGC

**Appendix Table S1. List of data measured for each strain, growth media and environment.**

| Media Envt. | | S^a^ | SEM | FOCI  rate^b^ | F_FOCIagg_^c^ | F_CYTOagg_^d^ | F_TOTALagg_^e^ | F_TOTALsol_^f^ | F_FOCIagg_ F_TOTALagg_ | F_CYTOagg_ F_TOTALsol_ | Coeff.^g^ |
| --- | --- | --- | --- | --- | --- | --- | --- | --- | --- | --- | --- |
| 5FOA | Sorbitol | 0.013 | ±6.29E-04 | 0.708 | 5042 | 9613 | 14655 | 51114 | 0.344 | 0.188 | 0.091 |
|  | Proline | 0.007 | ±2.82E-04 | 0.478 | 1703 | 7377 | 9080 | 27854 | 0.188 | 0.265 | 0.027 |
|  | NaCl | 0.006 | ±2.08E-04 | 0.308 | 1917 | 5791 | 7708 | 31935 | 0.249 | 0.181 | 0.019 |
|  | H_2_O_2_ | 0.028 | ±1.38E-03 | 0.681 | 2869 | 5567 | 8436 | 33762 | 0.340 | 0.165 | 0.156 |
|  | DTT | 0.022 | ±1.01E-03 | 0.705 | 3989 | 5183 | 9172 | 31855 | 0.435 | 0.163 | 0.151 |
|  | 37°C | - | - | 0.63 | - | 1411 | 1704 | 18688 | - | 0.075 | - |
|  | 30°C | 0.055 | ±2.71E-03 | 0.584 | 2893 | 5854 | 8746 | 31595 | 0.331 | 0.185 | 0.285 |
|  | 25°C | 0.002 | ±6.46E-05 | 0.884 | 5370 | 4766 | 10136 | 32962 | 0.530 | 0.145 | 0.061 |
| -URA | Sorbitol | -0.003 | ±-6.14E-05 | 0.676 | 2445 | 5584 | 8029 | 27919 | 0.305 | 0.2 | -0.013 |
|  | Proline | -0.004 | ±-2.08E-04 | 0.579 | 1186 | 4487 | 5673 | 9850 | 0.209 | 0.456 | -0.009 |
|  | NaCl | -0.004 | ±-1.76E-04 | 0.297 | 606 | 3398 | 4005 | 23828 | 0.151 | 0.143 | -0.029 |
|  | H_2_O_2_ | -0.010 | ±-5.23E-04 | 0.247 | 477 | 3471 | 3949 | 24096 | 0.121 | 0.144 | -0.073 |
|  | DTT | -0.007 | ±-2.86E-04 | 0.197 | 490 | 3296 | 3786 | 17186 | 0.129 | 0.192 | -0.034 |
|  | 37°C | -0.103 | ±-5.08E-03 | 0.689 | 2660 | 2990 | 5650 | 11737 | 0.471 | 0.071 | -0.403 |
|  | 30°C | -0.045 | ±-2.19E-03 | 0.477 | 691 | 3183 | 3874 | 18281 | 0.178 | 0.174 | -0.258 |
|  | 25°C | -0.01 | ±-4.65E-04 | 0.301 | 1107 | 4893 | 6001 | 16015 | 0.184 | 0.306 | -0.031 |
| +URA | Sorbitol | -0.005 | ±-2.36E-04 | 0.865 | 5773 | 6559 | 12332 | 44318 | 0.468 | 0.148 | -0.011 |
|  | Proline | -0.001 | ±4.26E-05 | 0.471 | 1323 | 4491 | 5814 | 24718 | 0.228 | 0.182 | -0.002 |
|  | NaCl | 0.003 | ±8.17E-05 | 0.707 | 2413 | 6462 | 8874 | 38575 | 0.272 | 0.168 | 0.01 |
|  | H_2_O_2_ | 0.002 | ±2.77E-05 | 0.71 | 2961 | 5291 | 8252 | 42580 | 0.359 | 0.124 | 0.007 |
|  | DTT | -0.001 | ±-1.63E-05 | 0.321 | 579 | 3022 | 3601 | 31586 | 0.161 | 0.096 | -0.007 |
|  | 37°C | 0.001 | ±2.56E-05 | 0.578 | 1360 | 2885 | 4245 | 9008 | 0.320 | 0.091 | 0.002 |
|  | 30°C | 0.002 | ±7.36E-05 | 0.733 | 2853 | 5908 | 8762 | 29966 | 0.326 | 0.197 | 0.006 |
|  | 25°C | -0.005 | ±-1.38E-04 | 0.886 | 4209 | 7049 | 11258 | 34907 | 0.374 | 0.202 | -0.012 |

a. Selection coefficient is calculated as
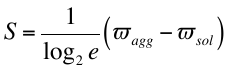
.

b. Ratio of cells with foci in that population.

c. Average fluorescence intensity emitted by a foci in the URA3_agg_ population. This is calculated as:

FOCIagg·FOCI_rate_

where FOCIagg is the average fluorescence intensity emitted by a foci taking into account only the fraction of URA3_agg_ population containing foci, and the FOCI_rate_ is the ratio of cells containing foci.

d. Average fluorescence intensity emitted by the cytoplasm of the URA3_agg_ population. This is calculated as: F_CYTOagg_ = F_TOTALagg_ – F_FOCIagg_

e. F_TOTALagg_ is the average of the total fluorescence emitted by URA3_agg_ population.

f. Average fluorescence intensity emitted by the URA3_sol_ population.

g. Coefficient of proportionality (i.e. magnitude of effect) defined by the growth conditions. It indicates how the growth conditions influences the magnitude of a particular effect (α, for deposit formation (+URA); β for loss of function (-URA); γ for gain of protective effect (+5FOA)).

**Appendix Table S2. List of doubling times measured for each strain, growth media and environment.**

|  | -URA (essential) | | +URA (non-essential) | | +5FOA (toxic) | |
| --- | --- | --- | --- | --- | --- | --- |
|  | URA3agg | URA3sol | URA3agg | URA3sol | URA3agg | URA3sol |
| 30ºC | 3.42±0.65 | 2.91±0.13 | 2.85±0.08 | 2.77±0.16 | 4.96±0.20 | 5.20±0.62 |
| Proline | 2.95±0.01 | 2.90±0.04 | 2.89±0.03 | 2.86±0.02 | 4.84±0.11 | 3.86±0.01 |
| NaCl | 3.24±0.05 | 3.17±0.02 | 3.15±0.01 | 3.12±0.09 | 5.39±0.20 | 4.76±0.07 |
| Sorbitol | 4.34±0.19 | 4.51±0.06 | 4.14±0.16 | 4.13±0.08 | 9.15±1.61 | 6.55±0.46 |
| H_2_O_2_ | 3.42±0.09 | 2.70±0.00 | 2.79±0.02 | 2.78±0.05 | 5.04±0.00 | 5.48±0.31 |
| DTT | 2.93±0.12 | 2.87±0.01 | 2.78±0.02 | 2.83±0.03 | 5.62±0.17 | 5.20±0.04 |
| 37ºC | - | 6.26±0.01 | 2.68±0.03 | 2.64±0.01 | 3.02±0.16 | 7.21±0.66 |
| 25ºC | 4.43±0.14 | 4.00±0.21 | 4.21±0.22 | 4.03±0.21 | 6.06±0.11 | 6.69±0.00 |

Doubling times calculated from the growth curve obtained by following the turbidity (OD600nm) and fluorescence (450 nm excitation & 510 nm emission) during 72 hours with a Tecan Infinite M200 Pro. Here we show the average doubling time value from two independent experiments. Time measured in hours.

**Appendix Table S3. Time required to reach 50% of the population in (simulated) alternating environments.**

| Environment 1 | | Environment 2 | | Selection coefficient of the history of environments (*S*_h_) | Time (in generations) to reach 50% of the population |
| --- | --- | --- | --- | --- | --- |
| H2O2 | FOA | 30 | FOA | 0.0415 | 126 |
| DTT | FOA | 30 | FOA | 0.0385 | 136 |
| Sorbitol | FOA | 30 | FOA | 0.0340 | 154 |
| Proline | FOA | 30 | FOA | 0.0310 | 169 |
| NaCl | FOA | 30 | FOA | 0.0305 | 171 |
| 30 | FOA | NaCl | URA+ | 0.0290 | 179 |
| 30 | FOA | 25 | FOA | 0.0285 | 185 |
| 30 | FOA | H2O2 | URA+ | 0.0285 | 185 |
| 30 | FOA | 30 | URA+ | 0.0285 | 185 |
| 30 | FOA | 37 | URA+ | 0.0280 | 187 |
| 30 | FOA | DTT | URA+ | 0.0270 | 195 |
| 30 | FOA | Proline | URA+ | 0.0270 | 195 |
| 30 | FOA | Sorbitol | URA- | 0.0260 | 203 |
| 30 | FOA | Proline | URA- | 0.0255 | 205 |
| 30 | FOA | NaCl | URA- | 0.0255 | 205 |
| 30 | FOA | Sorbitol | URA+ | 0.0250 | 212 |
| 30 | FOA | 25 | URA+ | 0.0250 | 212 |
| H2O2 | FOA | DTT | FOA | 0.0250 | 209 |
| 30 | FOA | DTT | URA- | 0.0240 | 216 |
| 30 | FOA | H2O2 | URA- | 0.0225 | 232 |
| 30 | FOA | 25 | URA- | 0.0225 | 232 |
| Sorbitol | FOA | H2O2 | FOA | 0.0205 | 255 |
| Proline | FOA | H2O2 | FOA | 0.0175 | 299 |
| Sorbitol | FOA | DTT | FOA | 0.0175 | 298 |
| NaCl | FOA | H2O2 | FOA | 0.0170 | 308 |
| H2O2 | FOA | NaCl | URA+ | 0.0155 | 339 |
| H2O2 | FOA | 25 | FOA | 0.0150 | 350 |
| H2O2 | FOA | H2O2 | URA+ | 0.0150 | 350 |
| H2O2 | FOA | 30 | URA+ | 0.0150 | 349 |
| Proline | FOA | DTT | FOA | 0.0145 | 360 |
| H2O2 | FOA | 37 | URA+ | 0.0145 | 360 |
| NaCl | FOA | DTT | FOA | 0.0140 | 375 |
| H2O2 | FOA | DTT | URA+ | 0.0135 | 387 |
| H2O2 | FOA | Proline | URA+ | 0.0135 | 387 |
| DTT | FOA | NaCl | URA+ | 0.0125 | 420 |
| H2O2 | FOA | Sorbitol | URA- | 0.0125 | 416 |
| H2O2 | FOA | Proline | URA- | 0.0120 | 439 |
| H2O2 | FOA | NaCl | URA- | 0.0120 | 439 |
| DTT | FOA | H2O2 | URA+ | 0.0120 | 437 |
| DTT | FOA | 25 | FOA | 0.0120 | 436 |
| DTT | FOA | 30 | URA+ | 0.0120 | 436 |
| DTT | FOA | 37 | URA+ | 0.0115 | 457 |
| H2O2 | FOA | Sorbitol | URA+ | 0.0115 | 455 |
| H2O2 | FOA | 25 | URA+ | 0.0115 | 455 |
| DTT | FOA | DTT | URA+ | 0.0105 | 497 |
| DTT | FOA | Proline | URA+ | 0.0105 | 497 |
| H2O2 | FOA | DTT | URA- | 0.0105 | 496 |
| Sorbitol | FOA | Proline | FOA | 0.0100 | 520 |
| DTT | FOA | Sorbitol | URA- | 0.0095 | 550 |
| Sorbitol | FOA | NaCl | FOA | 0.0095 | 547 |
| DTT | FOA | Proline | URA- | 0.0090 | 585 |
| DTT | FOA | NaCl | URA- | 0.0090 | 585 |
| H2O2 | FOA | H2O2 | URA- | 0.0090 | 579 |
| H2O2 | FOA | 25 | URA- | 0.0090 | 579 |
| DTT | FOA | Sorbitol | URA+ | 0.0085 | 614 |
| DTT | FOA | 25 | URA+ | 0.0085 | 614 |
| Sorbitol | FOA | NaCl | URA+ | 0.0080 | 649 |
| DTT | FOA | DTT | URA- | 0.0075 | 696 |
| Sorbitol | FOA | H2O2 | URA+ | 0.0075 | 693 |
| Sorbitol | FOA | 25 | FOA | 0.0075 | 692 |
| Sorbitol | FOA | 30 | URA+ | 0.0075 | 691 |
| Sorbitol | FOA | 37 | URA+ | 0.0070 | 740 |
| Proline | FOA | NaCl | FOA | 0.0065 | 805 |
| DTT | FOA | H2O2 | URA- | 0.0060 | 873 |
| DTT | FOA | 25 | URA- | 0.0060 | 869 |
| Sorbitol | FOA | DTT | URA+ | 0.0060 | 867 |
| Sorbitol | FOA | Proline | URA+ | 0.0060 | 860 |
| Proline | FOA | NaCl | URA+ | 0.0050 | 1046 |
| 30 | FOA | 30 | URA- | 0.0050 | 1029 |
| Sorbitol | FOA | Sorbitol | URA- | 0.0050 | 1024 |
| NaCl | FOA | NaCl | URA+ | 0.0045 | 1164 |
| Proline | FOA | H2O2 | URA+ | 0.0045 | 1163 |
| Proline | FOA | 25 | FOA | 0.0045 | 1162 |
| Proline | FOA | 30 | URA+ | 0.0045 | 1161 |
| Sorbitol | FOA | Proline | URA- | 0.0045 | 1150 |
| Sorbitol | FOA | NaCl | URA- | 0.0045 | 1150 |
| NaCl | FOA | H2O2 | URA+ | 0.0040 | 1311 |
| NaCl | FOA | 25 | FOA | 0.0040 | 1309 |
| NaCl | FOA | 30 | URA+ | 0.0040 | 1308 |
| Proline | FOA | 37 | URA+ | 0.0040 | 1305 |
| Sorbitol | FOA | Sorbitol | URA+ | 0.0040 | 1283 |
| Sorbitol | FOA | 25 | URA+ | 0.0040 | 1279 |
| NaCl | FOA | 37 | URA+ | 0.0035 | 1494 |
| Proline | FOA | DTT | URA+ | 0.0030 | 1752 |
| Proline | FOA | Proline | URA+ | 0.0030 | 1734 |
| Sorbitol | FOA | DTT | URA- | 0.0030 | 1693 |
| NaCl | FOA | DTT | URA+ | 0.0025 | 2113 |
| NaCl | URA+ | H2O2 | URA+ | 0.0025 | 2100 |
| 25 | FOA | NaCl | URA+ | 0.0025 | 2095 |
| NaCl | URA+ | 30 | URA+ | 0.0025 | 2093 |
| NaCl | FOA | Proline | URA+ | 0.0025 | 2086 |
| 25 | FOA | H2O2 | URA+ | 0.0020 | 2625 |
| H2O2 | URA+ | 30 | URA+ | 0.0020 | 2622 |
| 25 | FOA | 30 | URA+ | 0.0020 | 2614 |
| NaCl | URA+ | 37 | URA+ | 0.0020 | 2612 |
| Proline | FOA | Sorbitol | URA- | 0.0020 | 2578 |
| Proline | FOA | Proline | URA- | 0.0015 | 3537 |
| Proline | FOA | NaCl | URA- | 0.0015 | 3529 |
| H2O2 | URA+ | 37 | URA+ | 0.0015 | 3492 |
| 25 | FOA | 37 | URA+ | 0.0015 | 3477 |
| 37 | URA+ | 30 | URA+ | 0.0015 | 3472 |
| NaCl | FOA | Sorbitol | URA- | 0.0015 | 3436 |
| Sorbitol | FOA | H2O2 | URA- | 0.0015 | 3333 |
| Sorbitol | FOA | 25 | URA- | 0.0015 | 3320 |
| NaCl | FOA | Proline | URA- | 0.0010 | 5365 |
| NaCl | FOA | NaCl | URA- | 0.0010 | 5360 |
| NaCl | URA+ | DTT | URA+ | 0.0010 | 5351 |
| Proline | FOA | Sorbitol | URA+ | 0.0010 | 5197 |
| Proline | URA+ | NaCl | URA+ | 0.0010 | 5188 |
| Proline | FOA | 25 | URA+ | 0.0010 | 5133 |
| H2O2 | URA+ | DTT | URA+ | 0.0005 | 11065 |
| 25 | FOA | DTT | URA+ | 0.0005 | 10928 |
| DTT | URA+ | 30 | URA+ | 0.0005 | 10874 |
| NaCl | FOA | Sorbitol | URA+ | 0.0005 | 10443 |
| Proline | URA+ | H2O2 | URA+ | 0.0005 | 10400 |
| 25 | FOA | Proline | URA+ | 0.0005 | 10274 |
| Proline | URA+ | 30 | URA+ | 0.0005 | 10228 |
| NaCl | FOA | 25 | URA+ | 0.0005 | 10188 |

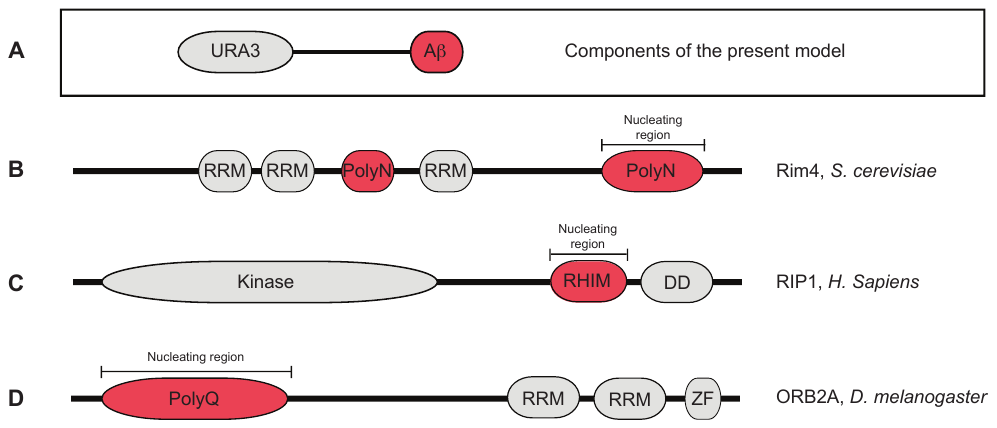

**Appendix Figure S1. Domain organization of proteins that can undergo phase separation and form functional deposits *in vivo*.**

**A.** The protein model for the current work has been designed to have a domain with a defined biochemical activity in the cell (enzymatic domain; URA3; grey) and a region that drives phase separation by simple coacervation (Aβ). Nature has selected for proteins with functions and properties separated into specific domains/regions. Here we present three natural proteins that phase separates by simple coacervation and contain spatially separated functional domains.

**B.** RIM4 represses several mRNAs capturing them in intracellular deposits. Rim4 aggregation is driven by a domain rich in Asparagines (polyN) and is independent of the presence of RNA presence (Berchowitz et al, 2015).

**C.** RIP1 and RIP3 are kinases that interact through a common region, the RIP homotypic interaction motif (RHIM). RHIM region drives the formation of amyloid deposits. *In vivo* RIP1 and RIP3 form a heterodimeric amyloid structure that mediates programmed necrosis (Li et al, 2012).

**D.** ORB2A is an isoform of ORB2, an orthologue of CPEB. ORB2A is crucial for the formation of ORB2 amyloid oligomers and the persistence of long-term memory (Khan et al, 2015).

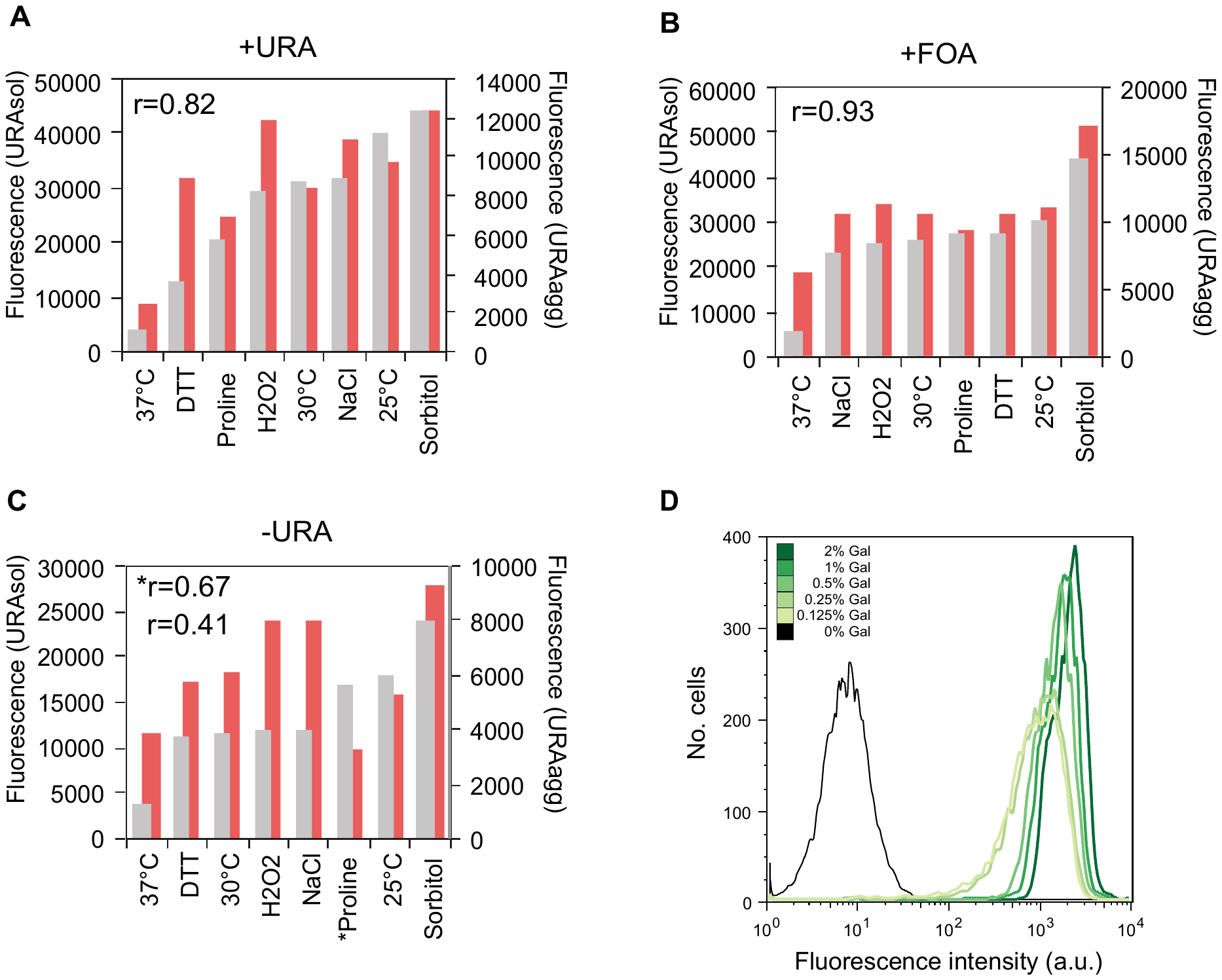

**Appendix Figure S2. Variation in fluorescence depending on the strain and the concentration of inducer.**

Fluorescence average (arbitrary units) of the URA3_agg_ (red, F_TOTALsol_) and URA3_sol_ (grey, F_TOTALagg_) populations grown at 30°C and in media with **A**, uracil, **B**, 5FOA or **C**, without uracil. Pearson’s correlation coefficient, *r*, is shown in each panel. Data is arranged from smaller to larger URA3_agg_ fluorescence values for each media composition. *Correlation coefficient obtained without the Proline environmental condition. Note: Although both strains (URA3_agg_ and URA3_sol_) present similar transcript levels (**Figure EV2**), due to deposit formation, a part of the translated Ura3p_agg_ can be rapidly removed through autophagy or other mechanisms (Sanchez de Groot et al, 2015; Villar-Pique & Ventura, 2013) and hence may lead to a reduction in F_TOTALagg_ with respect to F_TOTALsol_ (Sanchez de Groot et al, 2015; Villar-Pique & Ventura, 2013) (**Methods**).

**D.** Distribution of URA3_sol_ cells in terms of their total fluorescence after 20h of growth in different galactose concentrations. Each sample was centrifuged and suspended in PBS. The samples were vortexed for 1 min before measuring the fluorescence loss using a BD LSR II flow cytometer system (BD Biosciences). 10000 cells were counted at a maximum flow rate of 600 events per second. GFP fluorescence was measured using a 488 nm laser for excitation and a 525/50 nm band pass filter.

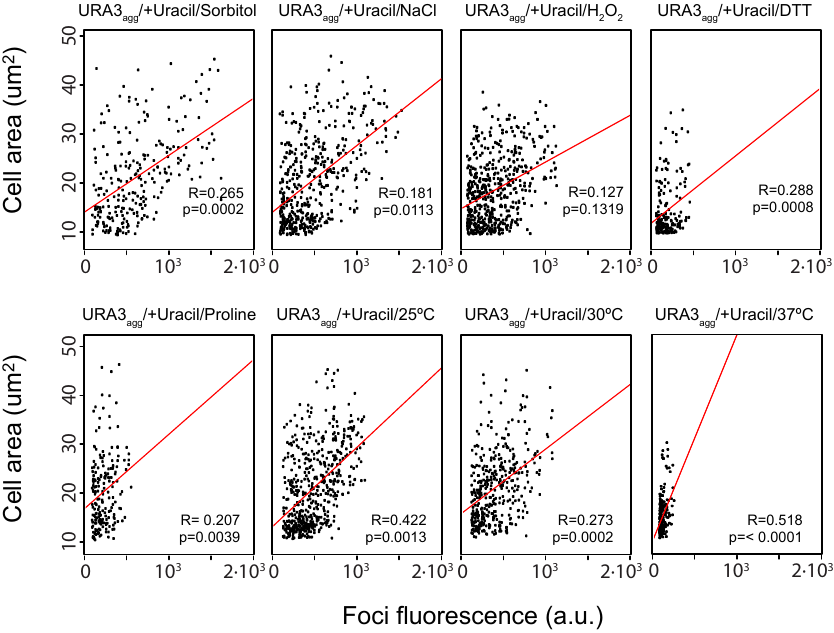

**Appendix Figure S3. Cell area vs. foci size distribution for URA3_agg_ cells grown in different environments when Ura3p is non-essential (+URA).** Scatter plots showing the relationship between the foci size and the cell area of the URA3agg population in a medium containing uracil (+URA). The environments are shown above. R denotes the Pearson’s correlation coefficient. P-values for the correlation coefficient were measured using the R statistical package.
