## Supplementary figures and images for "The fitness cost and benefit of phase separated protein deposits"

### Supplementary Figure EV1

**A**

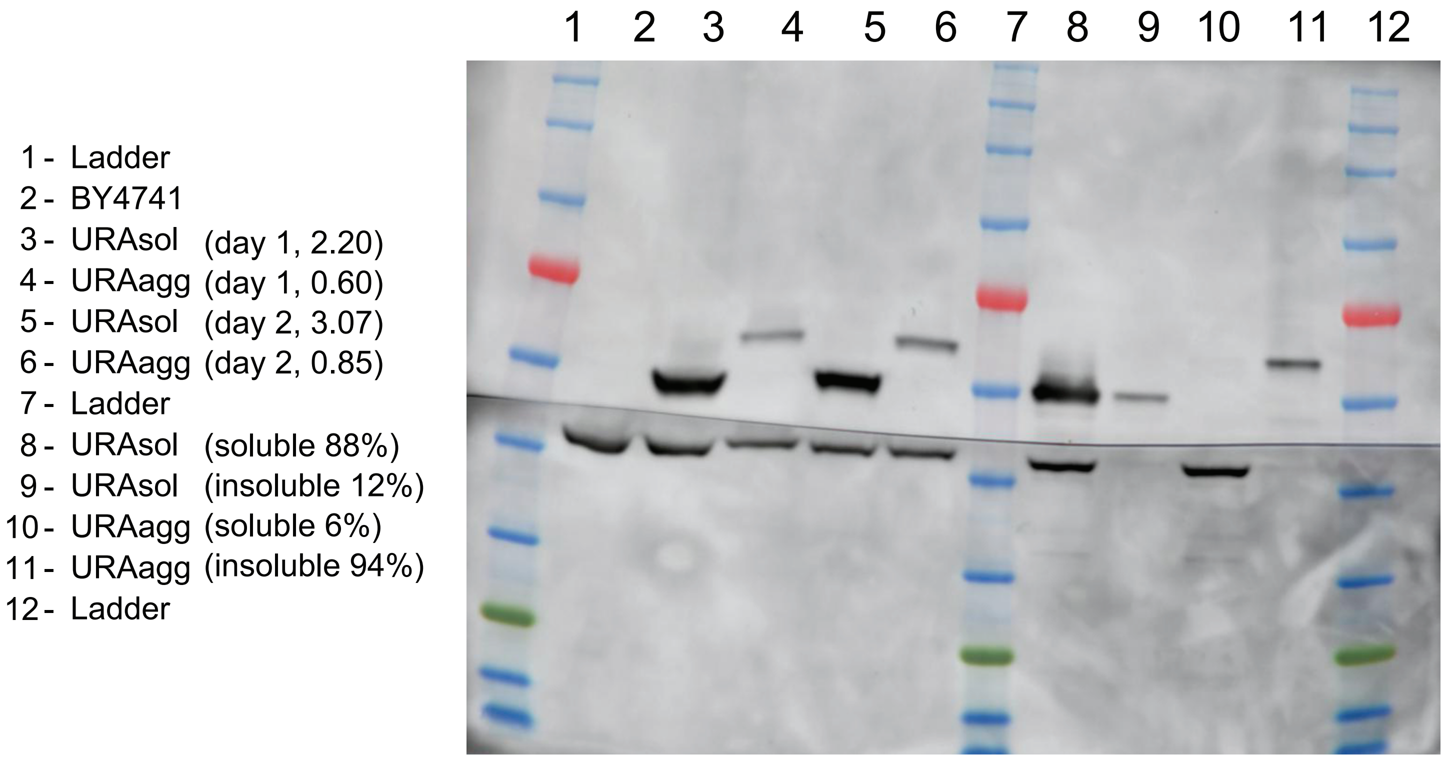


**B**

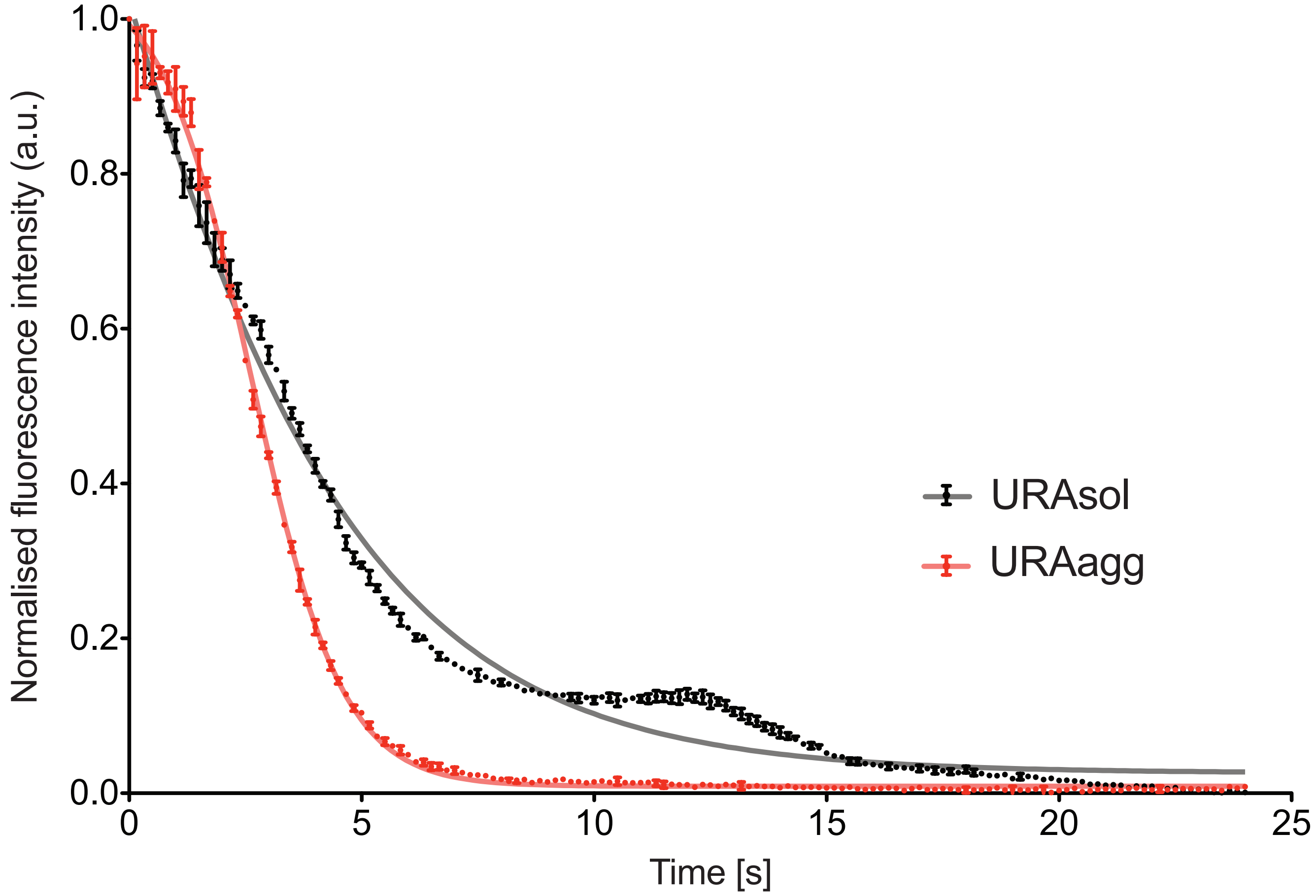


| | URAagg | URAsol |
|---|---|---|
| **Slope** | 3.456 | 0.9729 |
| **Std. Error** | 0.1192 | 0.0084 |
| $R^2$ | 0.9881 | 0.9987 |

### Supplementary Figure EV2

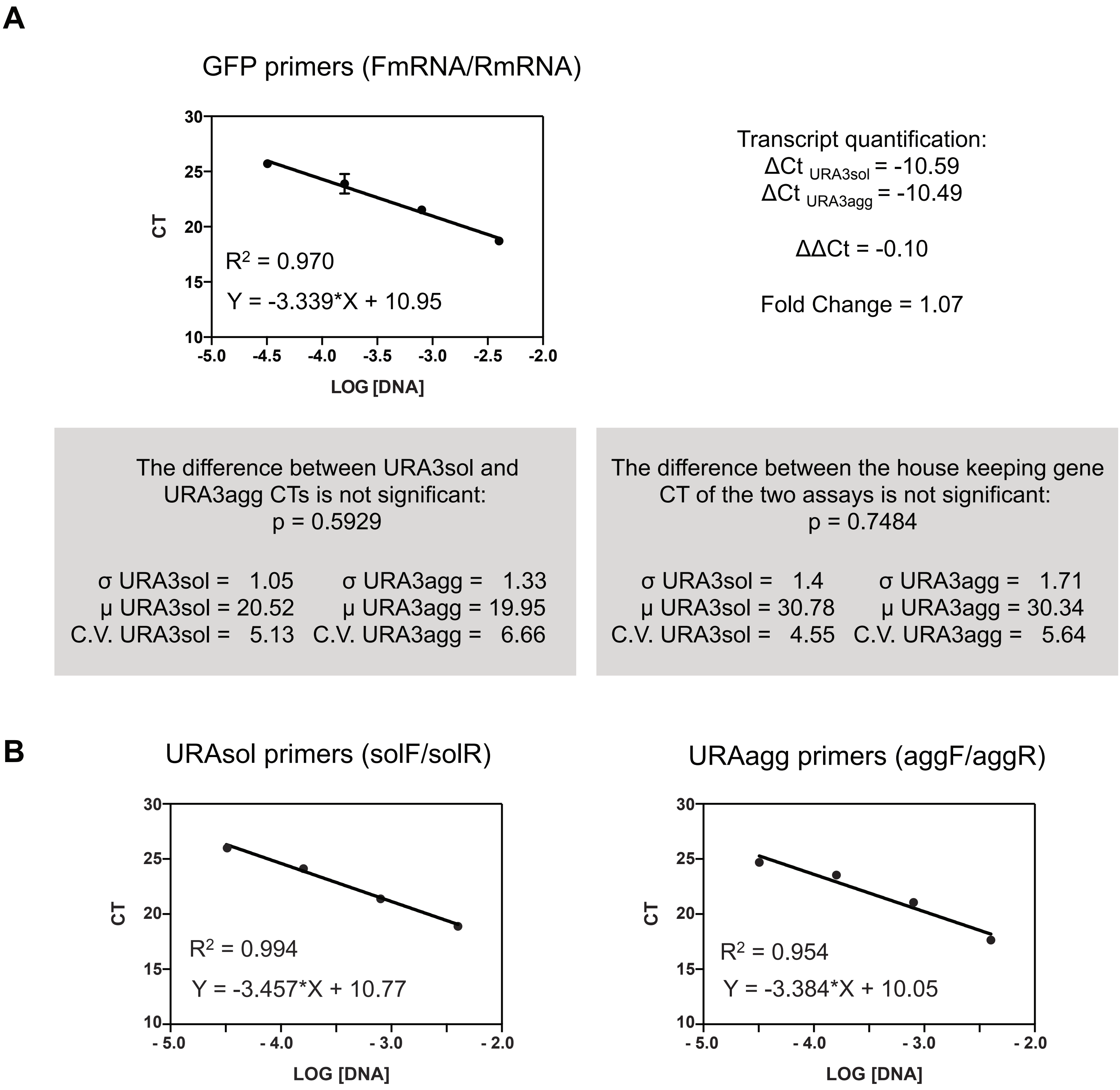

### Supplementary Figure EV3

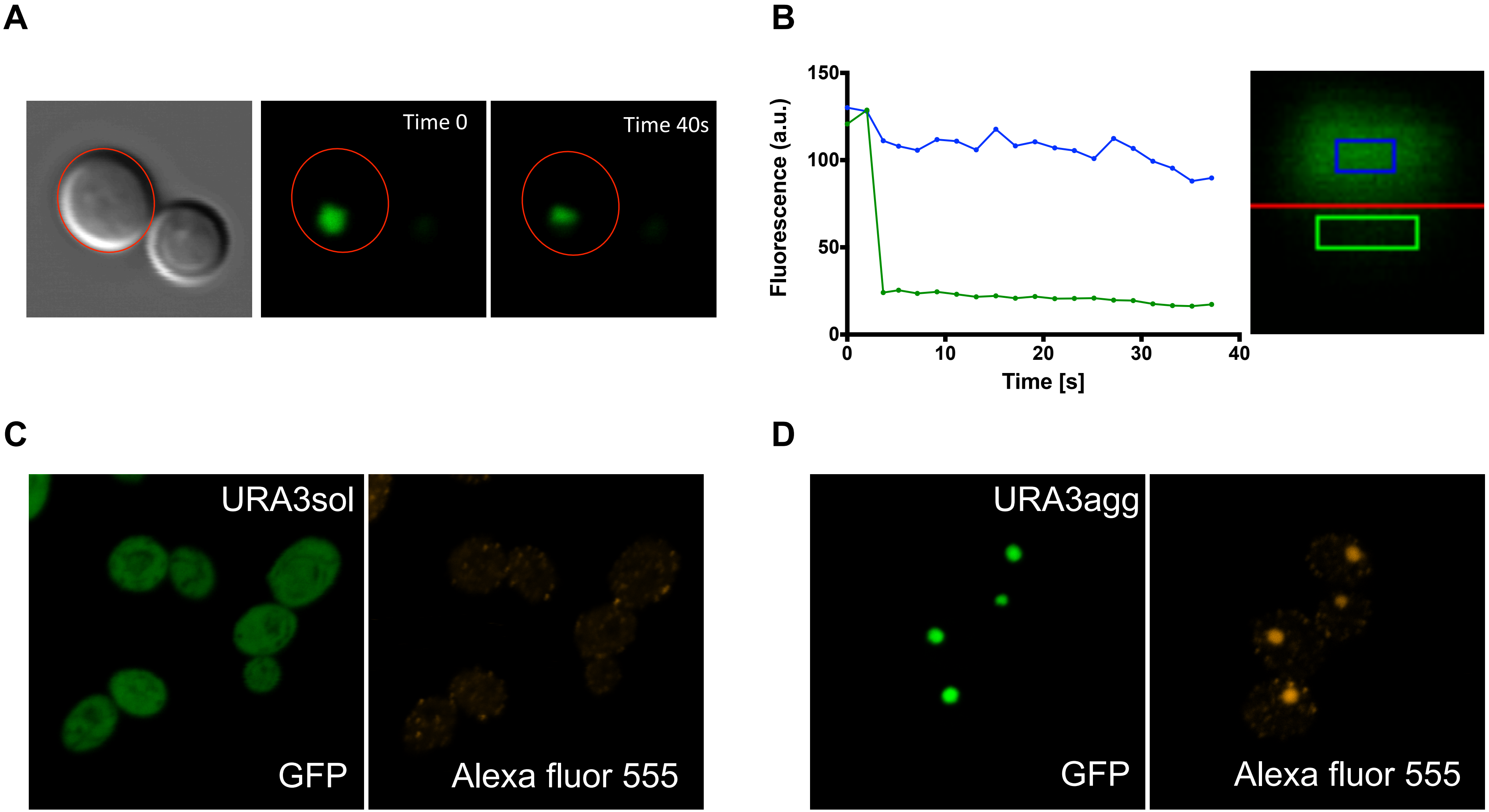

### Supplementary Figure EV5

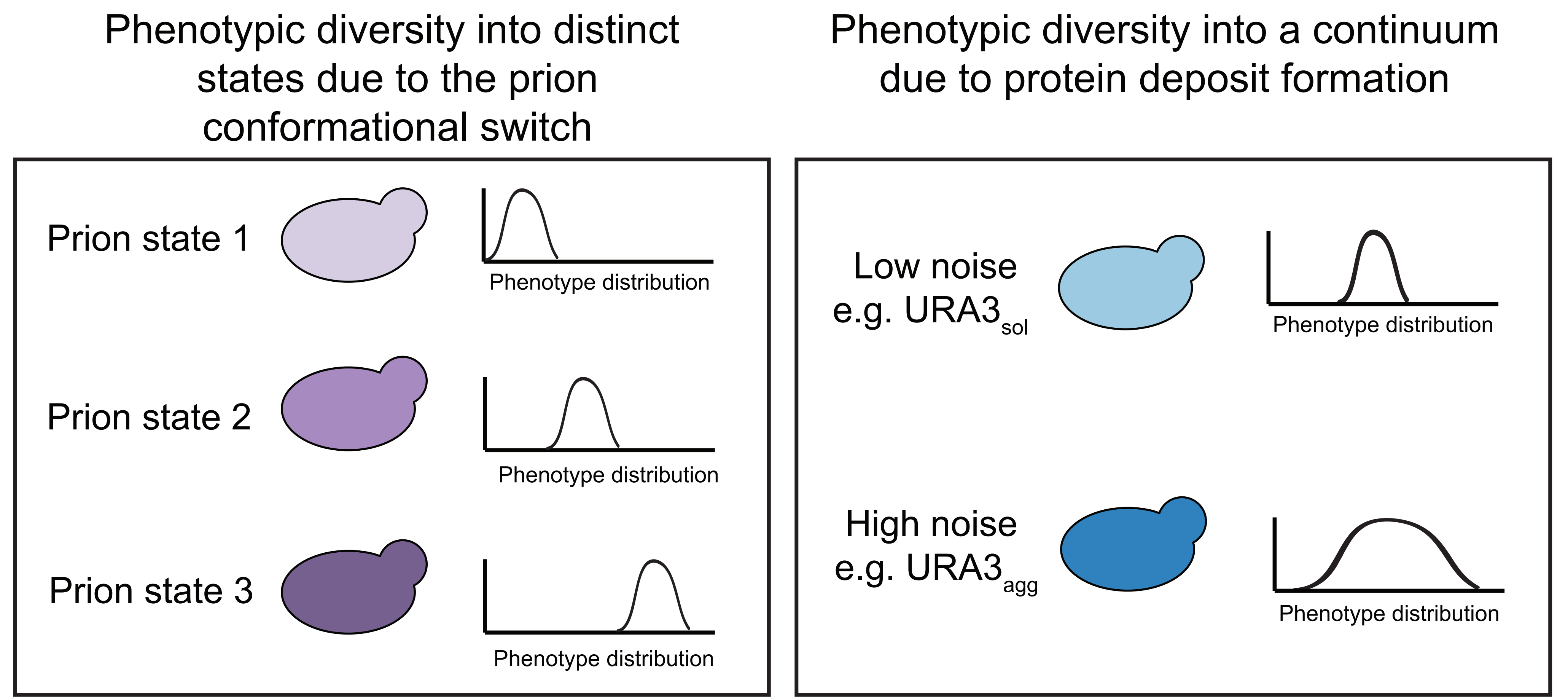
