## Supplementary Figure EV4 for "The fitness cost and benefit of phase separated protein deposits"

**A**

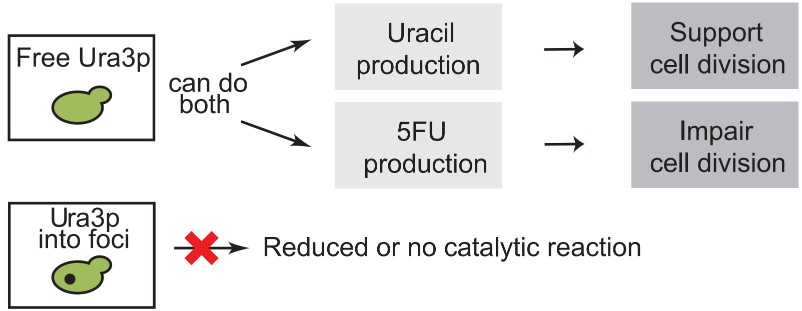


**B**

| ENVIRONMENT | ENVIRONMENT EFFECT | URA3p ACTIVITY | CELLULAR DEMAND | SELECTION OUTCOME | |
|---|---|---|---|---|---|
| Uracil in the media (+URA) | URA3p activity is not essential | Neutral | No demand | URA3sol ≈ URA3agg | Minor effects |
| Favours folding (25ºC, Pro, Sorbitol, NaCl) | Abundant free & active URA3p | Toxic<br>Essential | Demand met | URA3sol ≈ URA3agg<br>URA3sol ≈ URA3agg | Minor effects |
| Impairs folding (37ºC, $H_2O_2$, DTT) | Scarce free & active URA3p | Toxic<br>Essential | Demand not met | URA3sol << URA3agg<br>URA3sol >> URA3agg | Major effects |
| Optimal growth (30ºC) | Fast cell division faster uracil consumption | Toxic<br>Essential | Demand not met | URA3sol << URA3agg<br>URA3sol >> URA3agg | Major effects |

Foci size: small – large

Level of folded protein (GFP): low – high
